# Single-dose ethanol intoxication causes acute and lasting neuronal changes in the brain

**DOI:** 10.1101/2020.09.09.289256

**Authors:** Johannes Knabbe, Jil Protzmann, Niklas Schneider, Dominik Dannehl, Michael Berger, Shoupeng Wei, Christopher Strahle, Astha Jaiswal, Sophie Lugani, Hongwei Zheng, Marcus Krüger, Karl Rohr, Rainer Spanagel, Henrike Scholz, Ainhoa Bilbao, Maren Engelhardt, Sidney B. Cambridge

## Abstract

Alcohol intoxication at early ages is a risk factor for development of addictive behavior. To uncover neuronal molecular correlates of acute ethanol intoxication, we used stable-isotope labeled mice combined with quantitative mass spectrometry to screen over 2000 hippocampal proteins of which 72 changed synaptic abundance up to two-fold after ethanol exposure. Among those were mitochondrial proteins and proteins important for neuronal morphology, including MAP6 and Ankyrin-G. Based on these candidate proteins, we found acute and lasting molecular, cellular, and behavioral changes following a single intoxication in alcohol-naïve mice. Immunofluorescence analysis revealed a shortening of axon initial segments. Longitudinal two-photon *in vivo* imaging showed increased synaptic dynamics and mitochondrial trafficking in axons. Knockdown of mitochondrial trafficking in dopaminergic neurons abolished conditioned alcohol preference in *Drosophila*. This introduces mitochondrial trafficking as a process implicated in reward learning, and highlights the potential of high-resolution proteomics to identify cellular mechanisms relevant for addictive behavior.

## Introduction

First alcohol intoxication at an early age is a critical risk factor for later alcohol binging and the development of alcohol addiction (DeWit et al., 2000; Morean et al., 2014; Whelan et al., 2014). For example, college students who had their first acute alcohol intoxication prior to age 13 years were over three times more likely to develop alcohol addiction than those with an alcohol intoxication at 19 years or later (Henry et al., 2011; Hingson et al., 2003). Similar observations can be made in mice - a single exposure to an intoxicating ethanol dose increases alcohol consumption and alcohol relapse later in life (Camarini and Hodge, 2004; Fullgrabe et al., 2007). Hence, it is of critical importance in alcoholism research to understand the neurobiological consequences of a single alcohol intoxicating exposure.

One brain region that is especially susceptible to the intoxicating effect of alcohol exposure, in particular during peri-adolescence, is the hippocampus and behaviors mediated by the hippocampus have long been known to be sensitive to the acute effects of ethanol (White and Swartzwelder, 2004). Moreover, exposure to ethanol doses that produce intoxication cause consistent impairments in hippocampus-dependent learning and memory processes (Van Skike et al., 2019). For example, a single alcohol exposure can induce changes in synaptic plasticity (Wanat et al., 2009), as interference with long-term potentiation (LTP) in the hippocampus and alterations in spontaneous activity of pyramidal cells have been reported (White et al., 2000). Multiple studies also examined the effects of chronic ethanol exposure on the hippocampus and the brain which yielded valuable insights to our understanding of ethanol and the consequences of addiction. To investigate the development of addiction however, the identification of acute correlates of ethanol exposure hold great promise.

Here we set out to identify molecular alterations underlying cellular changes following acute ethanol intoxication and then investigated their influence on behavior. To achieve this, we focused on synapses as they are the main sites for neuronal transmission, plasticity, and information storage (Poo et al., 2016). We aimed to discover general principles of ethanol-induced changes and therefore performed a proteomic screen in the hippocampus. We choose the hippocampus as region of interest, because of its susceptibility to ethanol and its well described synapses. Acute *ex vivo* hippocampal slices display normal protein synthesis and degradation and are the standard model for plasticity studies (Fonseca et al., 2006). The hippocampus also shows less neuronal diversity (Sugino et al., 2019) and we expected that true, possibly minor stimulus-dependent protein abundance changes, are more discernable in homogenous samples of synapses. We conducted a proteomic screen in *ex vivo* slices from young mice (postnatal day - PND30) to capture the vulnerable peri-adolescent phase. Using slices, we aimed to reduce the inherent variability of living intoxicated animals.

Here, we found several dozens of proteins that significantly changed their synaptic abundance upon an acute ethanol stimulus. In our screen, we found a cluster of proteins indicating that mitochondria are crucial to mediate ethanol-dependent cellular changes, which we further tested. In addition, we identified microtubule-associated protein-6 (MAP6/STOP) and Ankyrin-G which suggests changes of neuronal morphology upon ethanol exposure. Based on these results, we next focused on morphological and mitochondrial dynamics in neurons *in vivo*. We found that the *ex vivo* results were robustly recapitulated in the brains of living mice as we could image and characterize changes in morphological and mitochondrial dynamics with two-photon microscopy. Intriguingly, the imaging data implied that mitochondrial trafficking and, in addition, lasting effects of ethanol could impact on ethanol-related behavior, which was separately verified in appropriate *Drosophila* and mouse behavioral tests, respectively.

## Results

### Identification of proteins that change their synaptic abundance upon acute ethanol exposure

To address whether a single ethanol intoxication is sufficient to cause structural and functional changes, we wanted to identify key protein contributors of *acute* ethanol intoxication, investigate their influence on relevant neuronal processes, and link these processes to behavior (**Fig. 1A**). To achieve this, we decided to investigate proteomes of synapses (Grant, 2018) as these are the most important structures that connect proteins to ethanol-related behavior. However, global, quantitative proteomic analyses of stimulus-dependent changes in synaptic protein abundance have been sternly limited by the heterogeneous nature of synapses, the potentially only subtle changes in synaptic abundance, and the incomplete fraction of synapses of a neuron that may actually respond to the stimulus. To overcome these limitations, we used quantitative high-resolution mass spectrometry (MS) in combination with acute hippocampal slices derived from ‘SILAC mice’ (SILAC: stable-isotope labeling in culture) (Kruger et al., 2008). With SILAC (Ong et al., 2003), labeled ‘heavy’ amino acids allow quantitative proteomics by measuring the peaks of the same peptides, from labeled and unlabeled samples, in one MS spectrum. For each protein, its relative abundance between samples is then calculated from the peptide peak intensities. In ‘SILAC mice’, all organismal lysines were metabolically replaced by stable-isotope labeled ‘heavy’ ^13^C_6_-lysine (+6 Da). SILAC ethanol-treated and WT untreated control hippocampal slices (or vice versa) were pooled in a cross-over design (**Supplementary Figure 1A**) prior to biochemical processing which eliminated any potential artefacts that might arise from differences in purification efficiencies between the different experimental conditions.

**Figure 1:**
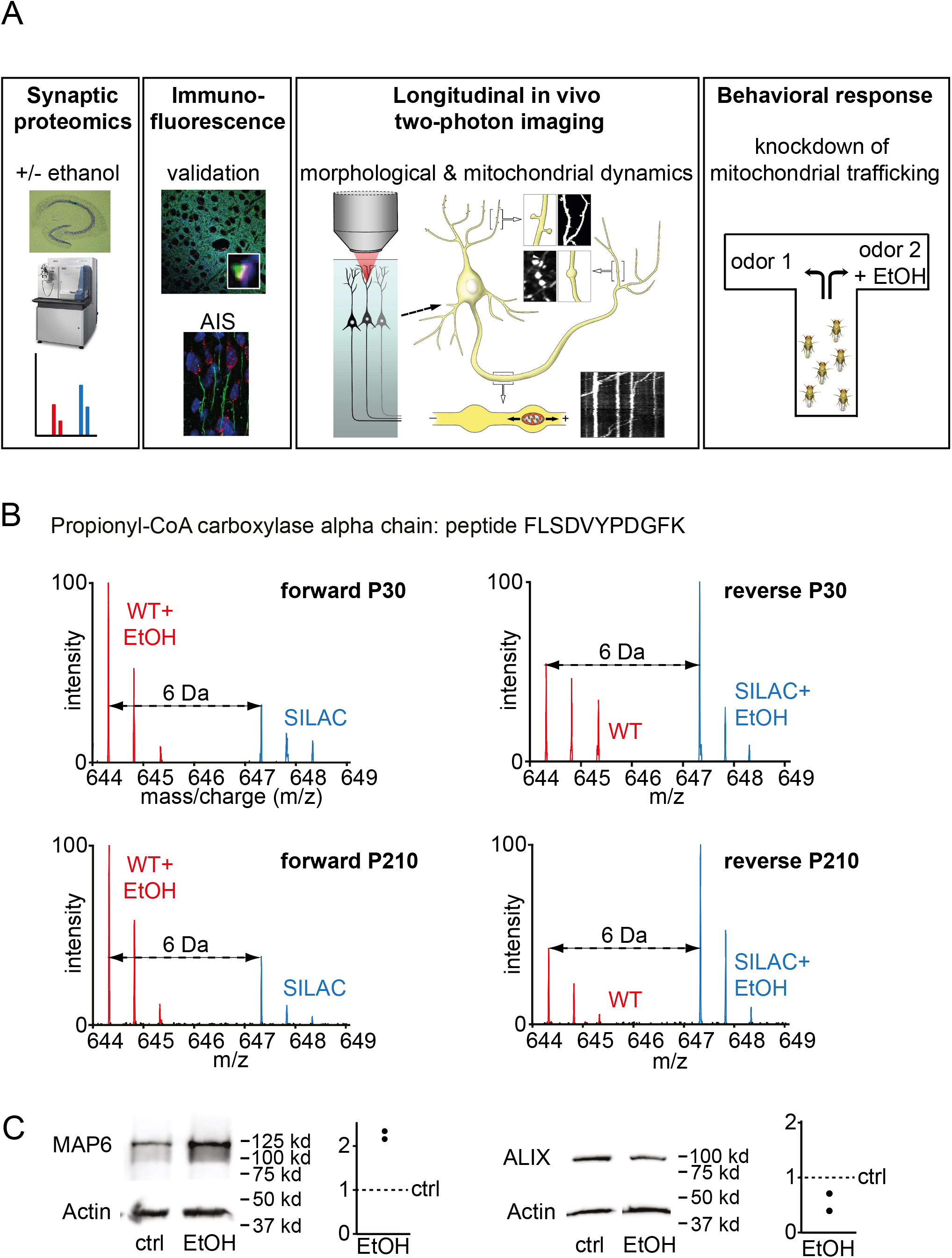
Study design and quantitative MS of synaptic proteomes. (**A**) Study design. (**B**) Raw MS spectra of WT and SILAC peptides. Reproducible and significant ethanol-dependent changes in peptide abundance of a peptide (FLSDVYPDGFK) unique to Propionyl-CoA carboxylase alpha. In all four experiments, ethanol exposure roughly doubled the peak height compared to untreated controls. (**C**) Western blot analysis of two candidate proteins. Actin was used as a loading control as MS results showed that it did not change between ethanol (EtOH) and control (ctrl) conditions. Numbers indicate molecular weight markers. Graphs show quantifications of Western blot experiments for candidate proteins by normalizing against actin controls.

We first established a small-scale experimental purification protocol for quantitative MS of highly purified synaptic preparations from acute slices derived from as little as two mouse hippocampi (**Supplementary Fig 1B, C**). We used Triton-X extraction at pH 6, because this not only retains the large, insoluble post-synaptic density (PSD) complex but also many presynaptic proteins (Phillips et al., 2001). We defined the proteins extracted by this enrichment procedure as ‘synaptic’. Applying our stringent requirements for MS identification on the data from the ethanol cross-over experiments produced a comprehensive, yet conservative estimate of 2089 pre- and postsynaptic proteins for the hippocampal synaptic proteome (**Supplementary Table 1**), including Homer, Bassoon, PSD-95, and many synaptic vesicle proteins. The number of detected proteins is in good agreement with previously published results (Pandya et al., 2017). In addition to the identification of synaptic proteins, our data also offer insight to the relative, i.e. stoichiometric abundance of each protein at the synapse (Wilhelm et al., 2014). Comparison of summed peptides intensities is a good proxy of the absolute protein abundance (Silva et al., 2006). Absolute quantification of 28 synaptic proteins (Cheng et al., 2006) showed that the abundance of 21 of these correlated well with our data set (**Supplementary Fig. 1D**).

For the cross-over experiment, a physiologically relevant and intoxicating ethanol concentration (4 h with 50 mM) was administered to acute *ex vivo* slices from young (PND30) mice and compared to control slices. The robustness of the *ex vivo* SILAC approach and quantitative analysis was demonstrated by the reproducibility of the ethanol-mediated changes at the peptide level. For each protein, the changes at the peptide-level were quantified with MaxQuant (Cox, 2008) to yield a relative, ethanol-dependent change of protein abundance at the synapse. As an example, **Figure 1B** shows the spectra of a unique peptide of the Propionyl-CoA carboxylase alpha chain (PCCA), a biotin-dependent mitochondrial enzyme, whose intensity increased similarly upon ethanol stimulation independent if labeled or unlabeled slices were used. Overall, in the screen we identified 57 proteins that significantly changed their level of abundance upon ethanol exposure (2.7% of 2089) (**Table 1**). We then conducted a second screen in acute hippocampal slices from adult mice (PND210) which strongly validated the first screen, as more than half of the changing proteins (32 out of 57) were detected again.

**Table 1:**
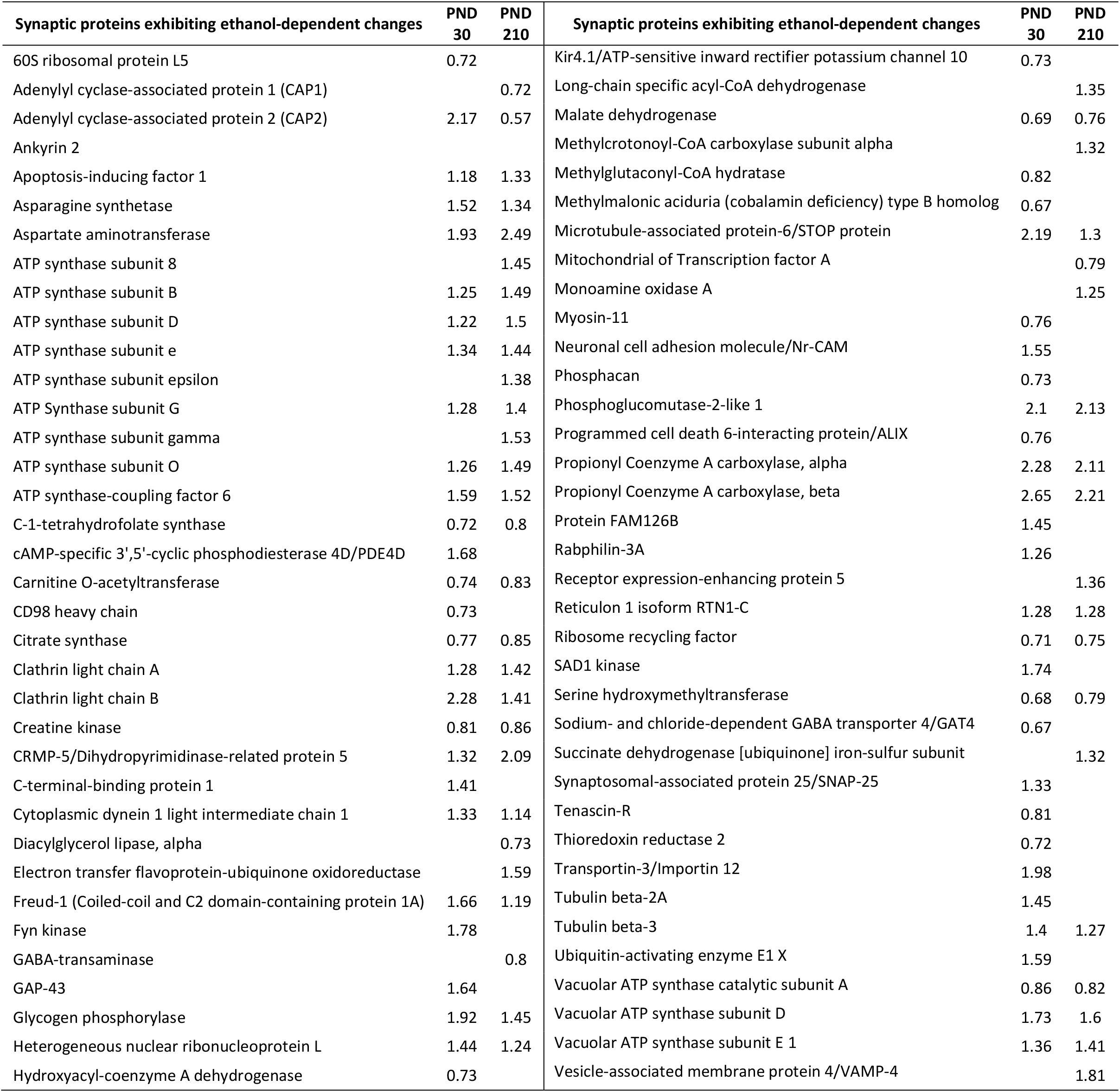
Proteins whose synaptic abundance changed significantly after ethanol. Ethanol-dependent changes in synaptic abundance for all significantly changing proteins. Ratios are given as fold change compared to controls. Depicted ratios are the average of forward and reverse experiments for young (PND30) or old (PND210).

| Synaptic proteins exhibiting ethanol-dependent changes | PND<br>30 | PND<br>210 | Synaptic proteins exhibiting ethanol-dependent changes | PND<br>30 | PND<br>210 |
| --- | --- | --- | --- | --- | --- |
| 60S ribosomal protein L5 | 0.72 |  | Kir4.1/ATP-sensitive inward rectifier potassium channel 10 | 0.73 |  |
| Adenylyl cyclase-associated protein 1 (CAP1) |  | 0.72 | Long-chain specific acyl-CoA dehydrogenase |  | 1.35 |
| Adenylyl cyclase-associated protein 2 (CAP2) | 2.17 | 0.57 | Malate dehydrogenase | 0.69 | 0.76 |
| Ankyrin 2 |  |  | Methylcrotonoyl-CoA carboxylase subunit alpha |  | 1.32 |
| Apoptosis-inducing factor 1 | 1.18 | 1.33 | Methylglutaconyl-CoA hydratase | 0.82 |  |
| Asparagine synthetase | 1.52 | 1.34 | Methylmalonic aciduria (cobalamin deficiency) type B homolog | 0.67 |  |
| Aspartate aminotransferase | 1.93 | 2.49 | Microtubule-associated protein-6/STOP protein | 2.19 | 1.3 |
| ATP synthase subunit 8 |  | 1.45 | Mitochondrial of Transcription factor A |  | 0.79 |
| ATP synthase subunit B | 1.25 | 1.49 | Monoamine oxidase A |  | 1.25 |
| ATP synthase subunit D | 1.22 | 1.5 | Myosin-11 | 0.76 |  |
| ATP synthase subunit e | 1.34 | 1.44 | Neuronal cell adhesion molecule/Nr-CAM | 1.55 |  |
| ATP synthase subunit epsilon |  | 1.38 | Phosphacan | 0.73 |  |
| ATP Synthase subunit G | 1.28 | 1.4 | Phosphoglucomutase-2-like 1 | 2.1 | 2.13 |
| ATP synthase subunit gamma |  | 1.53 | Programmed cell death 6-interacting protein/ALIX | 0.76 |  |
| ATP synthase subunit O | 1.26 | 1.49 | Propionyl Coenzyme A carboxylase, alpha | 2.28 | 2.11 |
| ATP synthase-coupling factor 6 | 1.59 | 1.52 | Propionyl Coenzyme A carboxylase, beta | 2.65 | 2.21 |
| C-1-tetrahydrofolate synthase | 0.72 | 0.8 | Protein FAM126B | 1.45 |  |
| cAMP-specific 3',5'-cyclic phosphodiesterase 4D/PDE4D | 1.68 |  | Rabphilin-3A | 1.26 |  |
| Carnitine O-acetyltransferase | 0.74 | 0.83 | Receptor expression-enhancing protein 5 |  | 1.36 |
| CD98 heavy chain | 0.73 |  | Reticulon 1 isoform RTN1-C | 1.28 | 1.28 |
| Citrate synthase | 0.77 | 0.85 | Ribosome recycling factor | 0.71 | 0.75 |
| Clathrin light chain A | 1.28 | 1.42 | SAD1 kinase | 1.74 |  |
| Clathrin light chain B | 2.28 | 1.41 | Serine hydroxymethyltransferase | 0.68 | 0.79 |
| Creatine kinase | 0.81 | 0.86 | Sodium- and chloride-dependent GABA transporter 4/GAT4 | 0.67 |  |
| CRMP-5/Dihydropyrimidinase-related protein 5 | 1.32 | 2.09 | Succinate dehydrogenase [ubiquinone] iron-sulfur subunit |  | 1.32 |
| C-terminal-binding protein 1 | 1.41 |  | Synaptosomal-associated protein 25/SNAP-25 | 1.33 |  |
| Cytoplasmic dynein 1 light intermediate chain 1 | 1.33 | 1.14 | Tenascin-R | 0.81 |  |
| Diacylglycerol lipase, alpha |  | 0.73 | Thioredoxin reductase 2 | 0.72 |  |
| Electron transfer flavoprotein-ubiquinone oxidoreductase |  | 1.59 | Transportin-3/Importin 12 | 1.98 |  |
| Freud-1 (Coiled-coil and C2 domain-containing protein 1A) | 1.66 | 1.19 | Tubulin beta-2A | 1.45 |  |
| Fyn kinase | 1.78 |  | Tubulin beta-3 | 1.4 | 1.27 |
| GABA-transaminase |  | 0.8 | Ubiquitin-activating enzyme E1 X | 1.59 |  |
| GAP-43 | 1.64 |  | Vacuolar ATP synthase catalytic subunit A | 0.86 | 0.82 |
| Glycogen phosphorylase | 1.92 | 1.45 | Vacuolar ATP synthase subunit D | 1.73 | 1.6 |
| Heterogeneous nuclear ribonucleoprotein L | 1.44 | 1.24 | Vacuolar ATP synthase subunit E 1 | 1.36 | 1.41 |
| Hydroxyacyl-coenzyme A dehydrogenase | 0.73 |  | Vesicle-associated membrane protein 4/VAMP-4 |  | 1.81 |

A total 72 proteins from PND30 and PND210 animals were identified whose synaptic abundance changed significantly upon the ethanol stimulus (**Table 1**). To confirm the ethanol-dependent changes, we then used Western blotting (WB) of synaptic proteins from acute hippocampal slices for one protein that showed an increase (MAP6), and for one protein that showed a decrease (ALIX or Programmed Cell Death 6-Interacting Protein) in synaptic abundance (**Fig. 1C**). The values obtained by Western blotting were in good agreement with the MS results for both proteins (MAP6↑: 2.2 [MS]/2.2 [WB]; ALIX↓: 0.76 [MS]/0.55[WB]). A Gene Ontology (GO) analysis was also conducted to reveal if certain molecular functions were particularly affected by acute ethanol intoxication. The enrichment of ATP synthase proteins was highly significant as were a few other molecular functions such as biotin binding, indicating alterations in mitochondrial functions (**Supplementary Fig. 2A**).

Among the 72 changing proteins, increases in synaptic abundance ranged from 1.25 to 2.65 fold-change and decreases from 0.85 to 0.57 fold-change. In total, 27 of the 72 candidate proteins (~38%) that changed significantly have been previously linked to ethanol (Ethanol-Related Gene Resource database, see (Guo et al., 2009)). Among these were proteins known to be affected by acute ethanol exposure such as malate dehydrogenase (Patel et al., 2000), monoamine oxidase A (Popova et al., 2000), GAP-43 (Kim et al., 2006), and Fyn tyrosine kinase (Miyakawa et al., 1997). For the latter protein, it has been shown that mice deficient for intracellular Fyn tyrosine kinase exhibited augmented alcohol intoxication (Miyakawa et al., 1997). Previous reports showed that ethanol augments GABA signaling by agonizing the GABA_A_ receptor (Faingold et al., 1998). Our MS data suggested that ethanol also reduced the synaptic abundance of the GABA re-uptake transporter GAT4 (**Supplementary Fig. 2B**) and GABA transaminase (Sherif et al., 1997), which catabolizes GABA. The reduction of both proteins should increase GABA signaling. An annotated, more detailed description of the identified proteins is presented in the Supplementary Material. Overall, the identified proteins link acute alcohol intoxication to alterations in synaptic transmission/plasticity, mitochondrial function, mood, apoptosis, and (neurodegenerative) diseases. In conclusion, the detected proteins support and expand literature knowledge and overlap with the Ethanol-Related Gene Resource, confirming the validity of our experimental approach.

We were intrigued to discover that Ankyrin-G and MAP6 were among the 72 candidate proteins because both have been reported to affect spine stability (Peris et al., 2018; Smith et al., 2014) while Ankyrin-G is also a key protein for the establishment and maintenance of the axon initial segment (AIS) (Leterrier, 2018). This suggested that acute ethanol exposure may induce detectable changes in neuronal morphology. Other notable candidate proteins included several mitochondrial proteins such as PCCA which indicated that the synaptic localization and trafficking of intact mitochondria was affected by the ethanol exposure. We therefore set out to visualize acute ethanol-dependent changes of neuronal morphology and mitochondrial trafficking in the brains of living mice. We reasoned that *in vivo* two-photon microscopy could be tailored to identify specific cellular changes in cortical structures of intoxicated animals.

### Synaptic protein abundance changes in the S1/M1 cortex *in vivo* after ethanol exposure

In preparation for the *in vivo* imaging experiments in the cortex, we verified that the protein candidates identified in hippocampal forebrain synapses also showed similar dynamics in cortical forebrain synapses. For this, we analyzed ethanol-dependent synaptic abundance changes in the primary somatosensory (S1)/motor (M1) cortex of intoxicated adult mice at different time points for up to 24 h after intraperitoneal (i.p.) injection of ethanol (3.5 g/kg body weight). Synaptic abundance of candidate proteins was quantified in brain sections by normalizing against presynaptic Synapsin and postsynaptic β-Actin, both of which did not significantly change upon ethanol exposure in the MS data set. We detected ethanol-induced synaptic accumulation of all three proteins that we analyzed, namely Ankyrin-G (**Fig. 2A-D**), MAP6 (**Fig. 2E-H**), and PCCA (**Fig. 2I-L**). Each protein significantly increased its synaptic abundance about 4 hours after injection (**Fig. 2M-O**) (P < 0.0001, Mann Whitney test; pooled 1-3 h after injection vs. 4-24 h after injection, 3 mice each) similar to the MS results (~1.5-2 fold) obtained with acute hippocampal slices.

**Figure 2:**
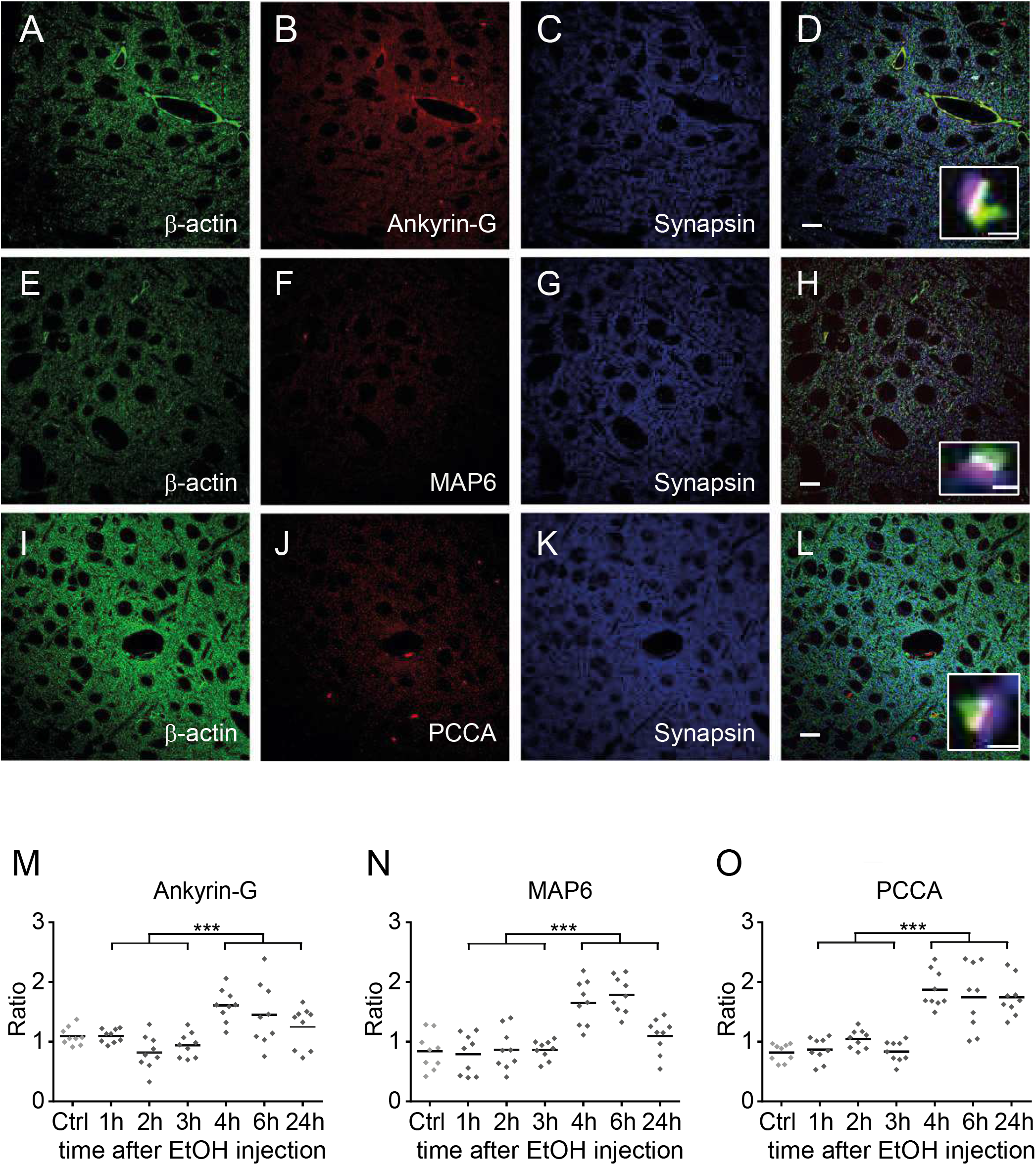
Immunofluorescence detection of ethanol-dependent synaptic protein dynamics *in vivo*. (**A-D**) Images for Ankyrin-G analysis, (**E-H**) images for MAP6 analysis, (**I-L**) images for PCCA analysis. (**B**) Ankyrin-G stain, (**F**) MAP6 stain (**J**) PCCA stain (**A, E, I**) ß-Actin stain. (**C, G, K**) Synapsin stain. Merged images (**D, H, L**). Insets: high magnification of synaptic labels (green: ß-actin, red: Ankyrin-G/MAP6/PCCA, blue; Synapsin). (**M**) Quantification of Ankyrin-G/Synapsin ratios over time, at 1, 2, 3, 4, 6, or 24 h after 3.5 g/kg ethanol i.p. injection. Each sample point corresponds to a single image as shown in this figure; each time point was based on one mouse. (**N**) Quantification of MAP6/Synapsin ratios over time. (**O**) Quantification of PCCA/Synapsin ratios over time. Scale bar A-L 20 μm, insets 1 μm.

Analysis of the persistence of synaptic accumulation also showed that the abundance of Ankyrin-G exhibited a more transient dynamic (**Fig. 2M**) compared to PCCA which remained elevated for at least 24 h after ethanol administration (**Fig. 2O**) (P = 0.0028, Mann Whitney test; pooled 4-24 h Ankyrin-G vs. PCCA, 3 mice) while MAP6 displayed an intermediate time course (**Fig. 2N**).We conclude that the results obtained from the proteomics screen with *ex vivo* acute hippocampal slices appear to adequately reflect ethanol-related changes at CNS synapses *in vivo*, at least for those proteins that we further characterized. Moreover, synapses exhibited substantial changes in protein composition after ethanol exposure *in vivo*, and these dynamics can be quite distinct for individual protein species.

### Ethanol-dependent morphological synaptic changes in dissociated cultures

We first wanted to study *in vitro* ethanol-induced spine dynamics in dissociated neuronal cultures. Acute exposure to 50 mM ethanol for 4 h induced a significant increase (~19 %) in spine density (P = 0.0002, Mann Whitney test, 3 experiments) (**Fig. 3A**, left). Interestingly, blocking neuronal activity with tetrodotoxin (TTX) resulted in a comparable increase in spine density (~13%) in cultures *in vitro* after 6 h (Papa and Segal, 1996). Since ethanol acts as a GABAmimetic via the GABA_A_ receptor (Forstera et al., 2016) and therefore is able to indirectly reduce neuronal activity, the observations suggest that ethanol might affect spine dynamics through activity-dependent mechanisms. Because the molecular composition of spines was dynamic in response to ethanol (**Fig. 2**), we next asked whether the morphological response to ethanol was in turn dependent on the molecular composition. Surprisingly, overexpression of Ankyrin-G (Smith et al., 2014) in combination with ethanol induced a significant decrease in spine density (P = 0.02, Mann Whitney test, 2 experiments) while overexpression of MAP6 (Peris et al., 2018) and ethanol had no effect (P = 0.85, Mann Whitney test, 2 experiments) (**Fig. 3A**, middle and right). Thus, the molecular composition of spines and dendrites appears to critically determine the morphological response to ethanol.

**Figure 3:**
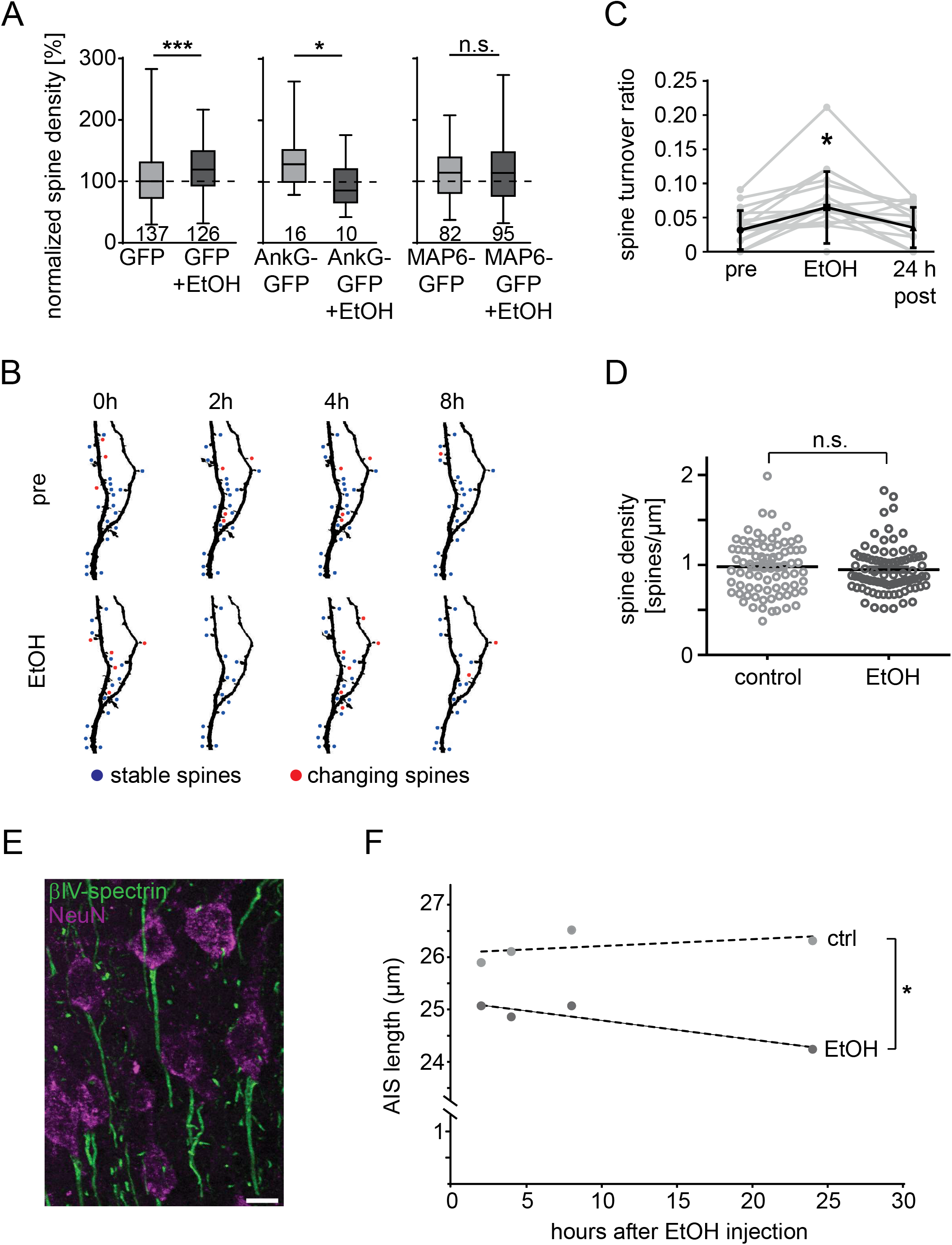
Neuronal *in vitro* and *in vivo* ethanol-dependent morphological plasticity. (**A**) Hippocampal cultures were transfected with Ankyrin-G-GFP, MAP6-GFP, or GFP at DIV8 and exposed to 50 mM ethanol for 4 h at DIV13. All Whisker plots were normalized to GFP control; numbers indicate dendritic stretches analysed. Ethanol alone induced a 19 % increase in spine density. Ethanol exposure in combination with Ankyrin-G expression decreased spine density but had no effect with MAP6. (**B**) Longitudinal *in vivo* two-photon imaging of the same dendritic stretches in Thy1-GFP mice identified stable (blue) and unstable spines (red). (**C**) Quantification of *in vivo* spine turnover from longitudinal imaging data (n= 4 mice; error bars: STD) (**D**) Ethanol intoxication in mice did not induce changes in cortical spine density after 4 - 6 h (6 mice for each condition) (**E**) Immunofluorescence detection of the AIS in layer II/III pyramidal neurons of S1 (green: βIV–spectrin, purple: NeuN) (scale bar: 10 μm). (**F**) Change in AIS length over time after a single ethanol i.p. injection (3.5 g/kg ethanol); about 100 AIS structures measured for each data point.

### Acute ethanol exposure causes changes in cortical spine dynamics *in vivo*

As a next step, spine dynamics were analyzed *in vivo* with longitudinal two-photon timelapse microscopy by imaging Thy1-GFP mice (Feng et al., 2000) in cortical layers II/III of M1/S1 cortex (**Fig. 3B**). Although we could not detect alterations in spine density at cortical synapses after ethanol administration (**Supplementary Fig. 2C**), there was a significant twofold increase in spine turnover after ethanol administration (**Fig. 3C**), which is indicative of increased structural plasticity. This was surprising since we observed an increase in spine density *in vitro* 4 h after the onset of ethanol exposure. To rule out that the rapid metabolism of ethanol *in vivo* diminished its effect on spine density, we blocked the activity of alcohol dehydrogenase by pyrazole in a different experiment and additionally kept the mice in ethanol vapor chambers for 4 h to maintain high blood ethanol concentrations (Eisenhardt et al., 2015). This *in vivo* exposure closely captured the ethanol conditions of the *in vitro* approach, but again we could not find any differences between control and ethanol-treated mice in spine density (**Fig. 3D**). In general, synapse dynamics *in vivo* are substantially lower than *in vitro* (Okabe, 2017) and a pronounced decline in postnatal *in vivo* spine formation was reported around PND21 (Isshiki et al., 2014). This could at least in part explain the ethanol-dependent increase in spine density we observed in two week old cultures, but not in adult, 2-4 month old mice.

We next addressed whether ethanol had an impact on the axon initial segment (AIS) as Ankyrin-G is a key protein for its establishment and maintenance (Leterrier, 2018). Modulation of both length and position of the AIS is an important mechanism for the regulation of intrinsic neuronal excitability in response to changes in network activity (Jamann et al., 2018). Quantification of AIS length in pyramidal neurons of layer II/III in S1 cortices (**Fig. 3E**) revealed a significant difference between ethanol-treated and control animals over time (P = 0.016, two-way repeated measures ANOVA) (**Fig. 3F**). Most notably, this minor, but significant effect of decreasing AIS length compared to controls continued up to 24 h, much beyond the actual presence of ethanol in the blood. This time course of AIS structural plasticity is in line with other reports showing changes in AIS length and diameter within hours and days after stimulus onset (Jamann et al., 2018).

### Increased mitochondrial mobility during and after acute ethanol intoxication

In our MS screen and GO analysis we found a cluster of proteins that suggested that intact mitochondria may play a role in mediating ethanol-dependent cellular changes. Most notable was the two-fold increase in PCCA that was seen by MS and *in vivo* immunofluorescence (**Fig. 2I-L**). We therefore decided to image and characterize the effects of ethanol on mitochondria in the cortex by *in vivo* two-photon time-lapse microscopy (**Supplementary Fig. 3A**). To be able to selectively monitor and analyze mitochondria in axons and boutons, two adeno-associated viruses (AAVs) with mitochondria-targeted GFP (mitoGFP) and a synaptophysin-mCherry (SyPhy-mCherry) as a marker for boutons were broadly injected into the thalamus (**Supplementary Fig. 3B**). As a consequence, fluorescence signal in the cortex (**Supplementary Fig. 3C**) derived from axonal projections of virally-transduced thalamic neurons. To assess the specificity of the effects of ethanol on mitochondrial transport compared to transport of other organelles, we also characterized dense-core vesicles (DCVs). DCVs were labeled via the cargo protein neuropeptide-Y (NPY) fused to the fluorescent protein Venus (NPY-Venus) under the same experimental *in vivo* conditions. We conducted longitudinal *in vivo* imaging of mitochondria and DCVs in addition to recording cortical 3Dstacks (“Z”) before, at 4 h, and at 24 h after i.p. injection of either saline or ethanol (3.5 g/kg) (**Fig. 4A**). Time-lapse movies of 10 min (“T”) were recorded in planes of layer I of M1, 50 - 100 μm away from the pial surface. The change in organelle trafficking was visualized in kymographs showing either idle, immobile mitochondria as straight lines over time (y-axis) or moving mitochondria with deviating paths (**Fig. 4B**). The two kymographs show identical regions before (top) and after (bottom) ethanol injection, which induced increased trafficking for a few mitochondria.

**Figure 4:**
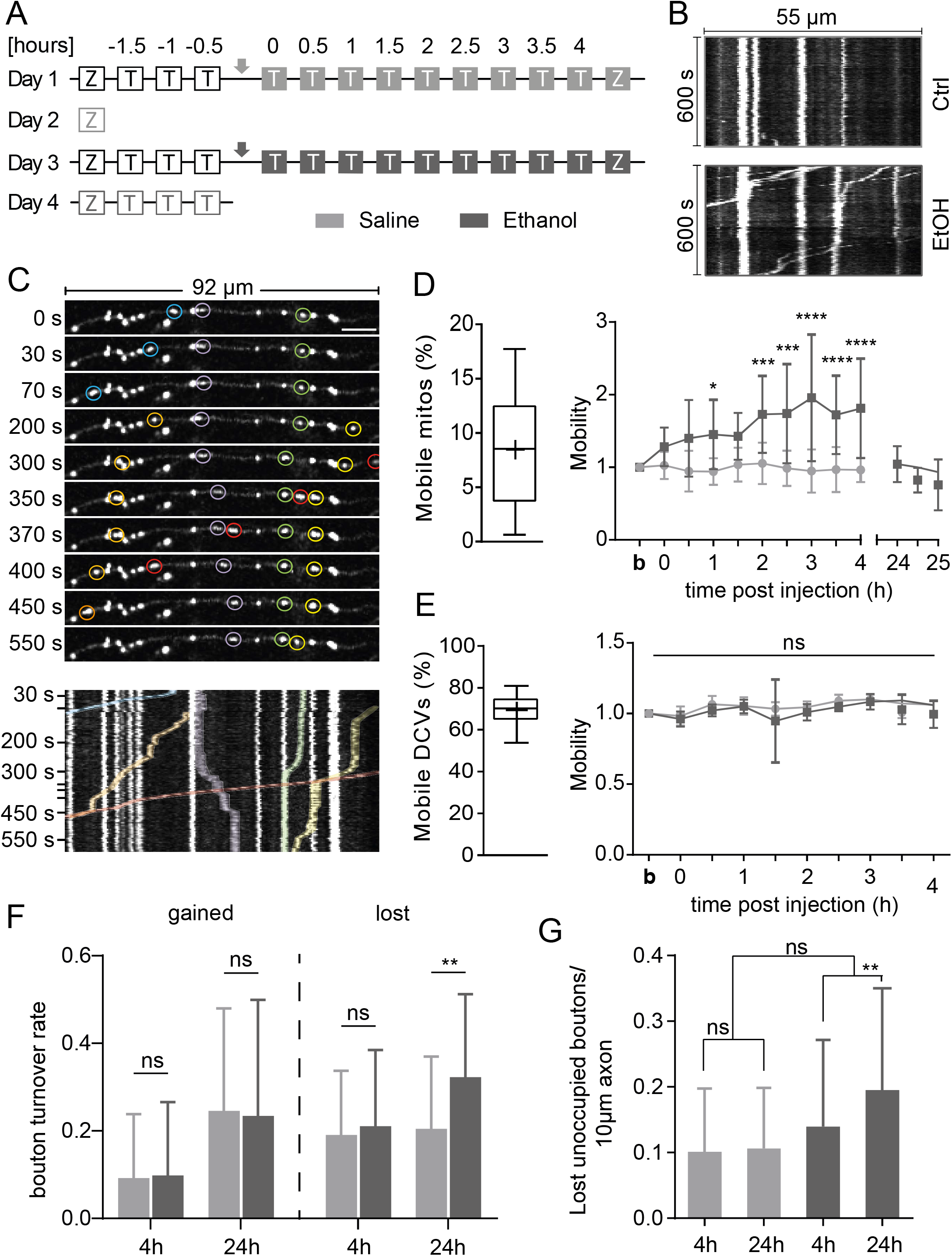
*In vivo* imaging of mitochondria and presynaptic boutons reveals ethanoldependent effects. (**A**) For each mouse, the experiment consisted of four imaging sessions. Baseline imaging (unfilled boxes) was acquired each day. On day 1, saline (light grey arrow) and on day 3 ethanol (dark grey arrow) were administered i.p. followed by post-injection imaging (filled boxes). On day 2 and 4, 24 h post-injection images were recorded. Z = 3D-Stack, T = time lapse. Time lapses (10 min mitochondria; 5 min DCVs) were acquired every 30 min. (**B**) Example kymograph before ethanol injection (top); only one mitochondrion was mobile. Kymograph of the same axonal stretch 180 min after ethanol injection (bottom). (**C**) Top: Axonal stretch at different time points (y-axis, seconds). Bottom: Corresponding kymograph. Orange: Slow mitochondrial trafficking. Red: Fast mitochondrial trafficking. Blue: Remobilization of a stable mitochondrion at 10 s. Purple: Relocation of a mitochondrion. Green: Fusion or co-localization of two mitochondria at 300 s. Yellow: Slow mitochondrial transport with intermittent pauses (100 – 200 s) and temporary fusion/co-localization (350 – 550 s) with another mitochondrion. White: Immobile mitochondria. (**D**) Left: Under basal conditions, 8.5 ± 4.7 % of mitochondria were mobile (n = 12 focal planes from 4 mice). Right: Time course of mitochondrial mobility under basal conditions (light grey) and after ethanol injection (dark grey). Mitochondrial mobility increased up to two-fold during acute ethanol intoxication, but returned to baseline (B) after 24 h (n =11 focal planes from 4 mice; mean ± STD). (**E**) Control. Left: Under basal conditions 69.4 ± 7.2 % of DCVs were mobile (n = 18 FOV from 3 mice). Right: Time course of DCV mobility under basal conditions (light grey) and after ethanol injection (dark grey). DCV mobility was not significantly different between saline and ethanol injected mice (n = 3 mice, mean ± STD). (**F**) Bouton turnover at different time points. 24 h after ethanol injection, increased bouton loss was observed compared to controls. (**G**) Loss of unoccupied presynaptic boutons was significantly increased after 4 h and 24 h after ethanol injection. Error bars: STD. (n = 40 axons from 4 mice). (Saline: light grey, Ethanol: dark grey)

**Figure 4C** shows an exemplary axonal stretch highlighting diverse aspects of axonal mitochondrial transport such as speed, direction of transport within the axon, and temporary pausing. First, mitochondrial transport was of variable speed: slow (orange) as well as fast (red) moving mitochondria were detected. Second, mitochondrial trafficking occurred in anterograde as well as in retrograde direction as organelles ‘move’ either left or right within the kymograph. In this recording, only the mitochondrion colored in purple moved into a different direction compared to all other highlighted mitochondria. Third, mitochondria alternated between stationary and mobile states, as a previously stationary mitochondrion started to move at approximately 10 s (blue). In general, we could detect a high number of stationary mitochondria that did not move during the time-lapse (white) (**Fig. 4C**, bottom).

Under baseline conditions *in vivo*, on average 8.47 ± 4.69 % of mitochondria were mobile (**Fig. 4D**, Supplementary Fig. 3D, **Supplementary Movie 1**). The mice then received an i.p. injection of saline or ethanol and we continued to image the same brain regions every 30 min for the following four hours. We found that over time, ethanol-dependent mitochondrial mobility increased compared to the saline controls (P = 0.0005, two-way repeated measures ANOVA, F (1,10) = 25.55) (**Fig. 4D**, **Supplementary Movie 2**). Acute ethanol intoxication significantly increased mitochondrial mobility after 30 min (P = 0.0152; Fisher’s uncorrected LSD test) and peaked at a two-fold increase after 180 min (P < 0.0001; Fishers LSD Test). This effect remained at least for 4 h but was not present anymore after 24 h. In contrast to mitochondrial mobility, mitochondrial velocity remained unchanged after acute ethanol intoxication (P = 0.4601, two-way repeated measures ANOVA) (**Supplementary Fig. 3E-G**).

To investigate whether the increase in mitochondrial transport mobility during ethanol exposure was specific for mitochondria or if it affected microtubule-based transport in general, we analyzed the movement characteristics of DCVs, which are predominantly transported via microtubules (Zahn et al., 2004) (**Supplementary Movie 3**). We found that on average 69.4 ± 7.16 % of DCVs were mobile under basal conditions (**Fig. 4E**) which is in good agreement with a previous study showing 75.1 % moving DCVs in axons of cultured mouse cortical neurons (de Wit et al., 2006). In contrast to mitochondrial transport, DCV transport mobility was not affected by ethanol intoxication (**Fig. 4E**). Similar to mitochondrial velocities, ethanol likewise did not increase DCV velocities (**Supplementary Fig. 3H-J**).

### Effects of higher mitochondrial mobility on occupancy of presynaptic boutons

Because of the increased disappearance/dislocation (**Supplementary Fig. 3K**) and higher mobility of mitochondria under ethanol, we next investigated whether this affected the dynamics of mitochondria occupancy within presynaptic boutons. Most stationary mitochondria are located at presynaptic terminals and are important during high synaptic energy demands (Verstreken et al., 2005) and synaptic plasticity (Smith et al., 2016; Vaccaro et al., 2017). Mitochondrial occupancy of synapses during ethanol intoxication was assessed by fluorescently labeling presynaptic boutons with virally transduced SyPhy-mCherry and quantification of co-localization with the mitoGFP signal (**Supplementary Fig. 4A**). Fluorescence co-localization was extracted from high-resolution Z-stacks (“Z”) at defined time points (**Fig. 4A**). We found that 80.5 ± 9.1 % of the presynaptic boutons on thalamic axons were occupied by mitochondria (**Supplementary Fig. 4B**) whereas other studies reported occupancy numbers between 42-60 % in axons in other neuron types and regions (Berthet et al., 2014; Lewis et al., 2016; Smit-Rigter et al., 2016). Four hours after ethanol injection we detected a small decrease in presynaptic mitochondrial occupancy (**Supplementary Fig. 4B**, right) which returned to baseline levels within 24h.

Analyzing the diameter of presynaptic boutons revealed that occupied stable boutons (1.25 ± 0.48 μm STD) were significantly larger (P < 0.0001, Kruskal-Wallis test with Dunn’s multiple comparison test) than unoccupied (0.89 ± 0.3 μm STD) or newly formed occupied boutons (0.91 ± 0.24 μm STD) (**Supplementary Fig. 4C**). We also analyzed mitochondrial shapes and found that ethanol did not affect mitochondrial length (**Supplementary Fig. 4D, E**).

Next, we wanted to see if we could detect ethanol-dependent morphological changes at presynaptic boutons. Bouton turnover was sub-grouped into newly gained (appeared during any of the imaging time points) or lost (present during baseline imaging that day but acutely disappeared) presynaptic boutons at different times points (4 h, 24 h) (**Fig. 4F**). Ethanol specifically and significantly increased the rate of bouton loss after 24 in the ethanol condition (P < 0.0001, two-way repeated measures ANOVA with Sidak’s correction for multiple comparisons; P = 0.0023 for interaction between condition and time; difference between ethanol and saline: P = 0.0508, F(1,78) = 3.936).

To further describe the heightened rates of bouton dynamics, presynaptic boutons with mitochondria were compared to unoccupied ones under ethanol conditions. Because we repeatedly imaged the same axonal stretches, we could clearly designate gained or lost boutons while also being able to identify boutons with gained or lost mitochondria (**Supplementary Fig. 5A-C**). In unoccupied boutons, ethanol significantly increased the rate of loss compared with the saline condition (P = 0.007, two-way repeated measures ANOVA, F (1,78) = 7.683) (**Fig. 4G**). This effect was significantly higher 24 h after ethanol compared to 4 h (P = 0.0011, Sidak’s multiple comparison test). Also, ethanol possibly affected the loss of occupied boutons (P = 0.0072 for difference between 4 and 24 hours in the ethanol condition; no difference between groups in a two-way repeated measures ANOVA with Sidak’s correction for multiple comparisons, F (1,78) = 0.05087), but not the formation of either unoccupied or occupied boutons (**Supplementary Fig. 5D-F**). Thus, ethanol exposure induced significant structural plasticity in presynaptic boutons and analogous to the AIS, lasting morphological changes existed at least up to 24 h after injection.

Because we observed lasting changes beyond 4 h and up to 24 h in form of molecular (PCCA) and cellular (AIS, boutons, mitochondria trafficking) correlates of acute ethanol intoxication, we wondered if these effects were mirrored by lasting changes in ethanol-related behaviors. For this purpose, we conducted a Go/NoGo task which is a measure for behavioral control and which is strongly affected by acute alcohol intoxication (Gubner et al., 2010). Naïve mice were first trained in a standard Go/NoGo task until they reached a designated performance level. The mice were then i.p. injected with 3.5 mg/kg ethanol and their performance assessed at 4 h, 24 h, and 48 h following injection. We found that at all time points after intoxication, mice exhibited significantly reduced performances in the correct go, false alarm, efficacy and performance rate (**Supplementary Fig. 6A-D**). Taken together, animals showed a post-intoxication effect of ethanol on behavioral inhibition, suggesting that one dose of ethanol is sufficient to cause longer lasting behavioral alterations.

### Mitochondrial trafficking is required to mediate positive rewarding properties of ethanol in dopaminergic neurons in *Drosophila*

The robust effects of ethanol on mitochondrial trafficking identified this cellular process as potentially relevant to regulate changes of neuronal plasticity underlying ethanol-induced behaviors. We therefore tested whether proper mitochondria trafficking is necessary to mediate the positive reinforcing effect of ethanol in dopaminergic reward neurons. We altered mitochondria trafficking in *Drosophila melanogaster* by knock-down of proteins required for axonal mitochondria transport and analyzed the consequences on ethanol-mediated reward learning and memory. Mitochondria are transported by the Miro/milton/kinesin complex to the synapse (Glater et al., 2006; Guo et al., 2005; Pilling et al., 2006; Stowers et al., 2002). In adult flies, a subset of dopaminergic neurons targeted by the *Tyrosine hydroxylase* (TH)-Gal4 driver mediates the positive rewarding properties of ethanol (Kaun et al., 2011). We expressed a RNA interference construct of *miro, UAS-dmiro-RNAi^TRiPJF02775^* under the control of the dopaminergic neuron specific *TH*-Gal4 driver. We then tested the experimental and the respective control groups in an olfactory associative learning and memory paradigm using ethanol as a positive reinforcer. A positive association between the olfactory stimuli and the unconditioned ethanol stimulus was achieved by exposing flies for 3 x 10 min to an odor cue that was reinforced with an intoxicating dose of ethanol vapor.

First, we verified that none of the olfactory stimuli were preferred and that the animals perceived the conditioned stimuli equally (**Supplementary Table 2**). Following the reinforcement conditioning, flies were given a choice in a T-maze between the rewarded odor cue and the non-rewarded odor cue. Consistent with previous results (Nunez et al., 2018), control conditioned animals significantly preferred the ethanol associated odor over the nonassociated odor cue. RNAi mediated knock-down of Miro in dopaminergic neurons however, significantly reduced the positive association between the olfactory stimulus and ethanol (**Figure 5A**).

**Figure 5:**
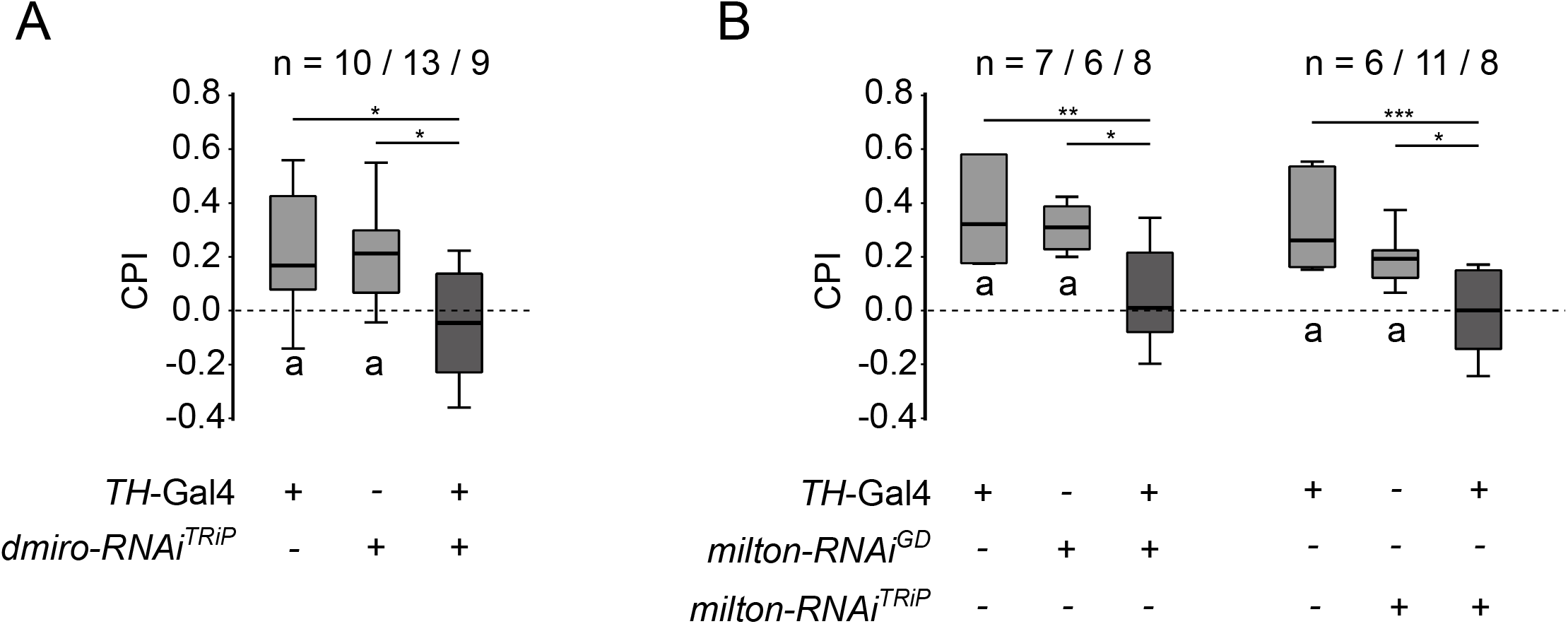
The function of ethanol as positive reinforcer in dopaminergic neurons depends on Miro and Milton. (**A**) The expression of the *UAS-dmiro-RNAi^TRiPJF02775^* under the control of the *TH*-Gal4 driver resulted in loss of CPI (conditioned odor preference). The mean of the CPI was for *TH*-Gal4/+: 0.22 ± 0.07; *UAS-dmiro-RNAi/+*: 0.21 ± 0.04 and for the experimental group −0.05 ± 0.07. (**B**) The expression of the *UAS-milton-RN4i^GD8116^* and *UAS-milton-RN4i^TRiPJF03022^* under the control of the *TH*-Gal4 driver resulted in loss of CPI. The mean of the CPI was for *TH*-Gal4/+: 0.37 ± 0.08, *UAS-milton-RNAi^GD8116^/+*: 0.31 ± 0.04 and for the experimental group 0.05 ± 0.06 and *TH*-Gal4/+: 0.32 ± 0.07; *UAS-milton-RN4i^TRiPJF03022^*/+: 0.19 ± 0.03 and for the experimental group −0.01 ± 0.06. The letter “**a**” indicates significant differences from random choice as determined with One-sample sign test. Differences between groups were determined using ANOVA *post-hoc* Tukey-Kramer HSD. Errors: SEM.

To confirm that the observed loss of reinforcement was specifically related to interference with mitochondrial trafficking, we also reduced Milton function with two different UAS-*milton*-RNAi lines, the UAS-*milton-RNAi^GD8116^* and the *UAS-milton-RNAi^TRiPJF03022^* (**Figure 5B**). Again, this RNAi-meditated knock-down in dopaminergic neurons resulted in a significant loss of the positive association with ethanol. We conclude that Milton/Miro-dependent mitochondrial trafficking is necessary for neuronal plasticity and behaviorally relevant positive rewarding properties of ethanol in dopaminergic neurons.

## Discussion

A single exposure of drugs of abuse including alcohol can have long-lasting behavioral consequences in rodents (Camarini and Hodge, 2004; Ciccocioppo et al., 2004; Halbout et al., 2014). In rodents, a single intoxicating exposure to ethanol increases alcohol consumption and alcohol relapse later in life (Camarini and Hodge, 2004; Fullgrabe et al., 2007). Here we studied the short-term and lasting molecular and cellular correlates of a single intoxicating exposure to ethanol. This was accomplished by investigating acute ethanol-dependent synaptic proteome changes, which revealed proteins that relate to mitochondrial function and neuronal morphology including the two structural proteins MAP6 and Ankyrin-G, that are important for spine stability and formation of the AIS. Thus, we show that high accuracy detection of minute, stimulus-dependent changes of synaptic protein abundance is now possible. To date, only a handful of proteins have been identified whose synaptic abundance reproducibly increased or decreased following any stimulus, including LTP (Hayashi et al., 2000; Strack et al., 1997). In our quantitative screen, PCC alpha and beta chains (PCCA/B) for example displayed two of the most marked increases (> two-fold). This protein has not been reported in the literature in an ethanol-related context. Because of the strong and reproducible ethanol-dependent increase, we envision that PCCA/B could be candidates to be developed into markers for acute and chronic alcohol intake. A similar increase in synaptic abundance was seen for aspartate aminotransferase, a protein mostly reported for chronic ethanol exposure and a known marker for alcoholism (Conigrave et al., 2003).

We then demonstrated with longitudinal two-photon time-lapse microscopy *in vivo* (i) a two-fold increase in spine turnover and increased bouton elimination after acute ethanol administration indicative of increased structural plasticity and (ii) a pronounced increase in mitochondrial mobility after acute ethanol intoxication. In addition, we found (iii) a lasting decrease in AIS length. The observed changes in mitochondrial trafficking were indeed of behavioral relevance. Genetic interference with mitochondrial trafficking within the dopaminergic reward pathway was required to mediate the positive reinforcement properties of ethanol. In conclusion, we introduce enhanced mitochondrial mobility and dynamics in presynaptic boutons as a key cellular mechanism which underlies alcohol reward learning and which is possibly a general cellular process important for learning and memory.

In neurons, there is a vast body of knowledge describing lasting morphological changes, including plasticity-related synaptic changes following a LTP or LTD stimulus (Engert and Bonhoeffer, 1999; Nagerl et al., 2004) as well as long-term adaptive morphological changes as consequence of neuronal homeostasis (Turrigiano, 2012; Yin and Yuan, 2014). We show here lasting, possibly homeostatic morphological changes following a single ethanol stimulus. Ethanol treated animals exhibited a significantly reduced AIS length compared to controls, a decrease which persisted at least up to 24 h after ethanol injection. Heightened neuronal activity has been reported to decrease AIS length (Evans et al., 2015). One conceivable interpretation of our data is based on published observations showing that acute ethanol rapidly decreases cortical activity (Lebedeva et al., 2017; Tu et al., 2007). Consequently, the lasting homeostatic AIS changes are possibly reflective of an ensuing network hyperactivity, because ethanol was rapidly metabolized following the initial ethanol-dependent silencing of neurons. Such a scenario would have to be confirmed by the difficult task of carefully quantifying neuronal activity at different time points *in vivo* following an ethanol stimulus. Besides the AIS, synapses also showed ethanol-dependent structural remodeling as changes in bouton turnover lasted up to 24 h. Albeit small, the observed loss of boutons could be interpreted as an adaptive homeostatic response as well. Intriguingly, homeostatic mechanisms have been reported for chronic ethanol exposure (Carpenter-Hyland and Chandler, 2006; Spanagel and Kiefer, 2008)

Morphological remodelling of neurons is a known substrate for learning and memory, as increased spine and bouton turnover has been found to be associated with memory formation (Ash et al., 2018; Frank et al., 2018; Roberts et al., 2010). Cellular plasticity mechanisms that are central to learning and memory are likewise considered to be at the core of forming associative memories for drug-related rewards (Hyman et al., 2006). Thus, some of the observed ethanol-dependent morphological changes we detected, i.e. the increased spine turnover and the elimination of boutons, could potentially influence ethanol-related memory formation by distorting the synaptic connectivity balance between spines and boutons. Together with trafficking of mitochondria, which are known to be important for synaptic transmission and plasticity (Todorova and Blokland, 2017), one could therefore speculate that these ethanol-dependent cellular changes are crucial substrates for the development of addictive behavior. We tested this idea by specifically blocking mitochondrial trafficking in a well-described model of ethanol reward in adult *Drosophila*. Similar to mice and humans, the formation of positive memories associated with ethanol intake is dependent on dopaminergic neurons in the *Drosophila* model. Knockdown of either known adaptor proteins, Miro and Milton, eliminated the preference for odor cues associated with ethanol intoxication.

Both mechanisms, i.e. long-lasting homeostatic structural remodeling and mitochondrial trafficking, may as well account for the observations made in mice that a single exposure to an intoxicating dose of ethanol can increase alcohol consumption and alcohol relapse later in life (Camarini and Hodge, 2004; Fullgrabe et al., 2007). The mechanisms may even be of relevance for the observation in humans that an early age of first alcohol intoxication is a critical risk factor for later alcohol binging and the development of alcohol addiction (DeWit et al., 2000; Morean et al., 2014; Whelan et al., 2014).

## Acknowledgements

We thank Gabriele Krämer, Marion Schmitt, Claudia Koksch, Michaela Kaiser for technical assistance, Christine Opfermann-Rüngeler for help with illustration. We are indebted to Thomas Kuner for continuous support of the project and comments on the manuscript.

## Funding

The project was in part funded by a grant of the Volkswagen-Stiftung to SBC, by a DFG-SCHO10/1 grant to HS, by the Promotion Fellowship from the Medical Faculty Mannheim, Heidelberg University for DD. Financial support for AB and RS was provided by the Bundesministerium für Bildung und Forschung (BMBF) funded SysMedSUDs consortium (FKZ: 01ZX1909A), and the Deutsche Forschungsgemeinschaft (DFG, German Research Foundation) – Project-ID 402170461 – TRR 265 (Heinz et al., 2020).

## Author Contributions

SC conceived and designed the study, wrote the manuscript. JK, HS, AB, ME designed and analyzed experiments, contributed to writing the manuscript. JP, NS, DD, MB, SW, CS, SL, SC performed and analyzed experiments. AJ and KR developed the automated tracking algorithm and analyzed data. HZ contributed to the spine analyses, MK to the MS experiments. RS contributed to writing the manuscript. All authors reviewed and approved the manuscript.

## Declaration of interest

The authors declare no competing interests

## Reagent and Data availability

Upon reasonable request, all unique/stable reagents and codes generated for mitochondrial tracking and spine analyses are available via the Lead Contact. The raw mass spectrometry data can be accessed at https://heidata.uni-heidelberg.de/privateurl.xhtml?token=f88c8325-6f37-473d-856a-0af8109a2bba.

Further information and requests for resources and reagents should be directed to and will be fulfilled by the Lead Contact Sidney Cambridge

## Supplementary Material

### Supplementary Figure Legends

**Supplementary Figure 1: Proteomic analyses of synaptic proteins.**

**A.** Cross-over experimental scheme.

**B.** Multi-step, small scale synapse purification procedure (P = pellet, S = supernatant).

**C.** Western blot analysis of the purification procedure. The Triton-X insoluble PSD-95 protein was stepwise enriched; the Triton-X soluble Synaptophysin protein was removed during the repeated Triton-X extractions. The same amount of total protein was loaded onto each lane (hom = homogenate, syn = synaptosomes)

**D.** Comparison of published AQUA data of postsynaptic density proteins with MS peak intensities obtained in this study. Cheng et al. (Cheng et al., 2006) determined absolute concentrations of 28 proteins in forebrain PSD preparations by AQUA (absolute protein quantification). 21 of these proteins (orange triangles) are in good agreement with the average peak intensities of our MS analysis of the pre- and postsynaptic hippocampal proteome (blue squares). Those 7 proteins with more than one magnitude difference between the AQUA and our MS data are highlighted in yellow and annotated.

**Supplementary Figure 2: GO analysis, GAT4 spectra, and *in vivo* spine densities**

**A**. Gene Ontology enrichment analysis of the 72 significantly changing proteins vs. detected hippocampal synaptic proteome. The numbers indicate: significantly changing protein of GO term / total number of GO term proteins detected in synaptic proteome (total number of GO term proteins in genome).

**B.** MS spectra of a unique peptide (LTVPSADLK) of the GABA transporter 4 protein. In both forward and reverse ethanol treated samples (P30), there was a significant reduction of the peptide/protein.

**C**. The distribution of spine densities during longitudinal *in vivo* 2-Photon imaging of Thy1-GFP mice revealed no significant changes before, during, or after ethanol exposure. (spine density in spines/μm: day 1: baseline - pre Ethanol: 0.8525, day 2: Ethanol: 0.8514, day 3: post Ethanol: 0.8549; repeated measures ANOVA p = 0.0001; Bonferroni’s multiple comparisons test p > 0.44)

**Supplementary Figure 3: Visualizing axonal organelle trafficking under saline and ethanol conditions using *in vivo* two-photon microscopy.**

**A**. Two-photon imaging was accomplished by implanting a chronic cranial window. The 3D-printed crown, needed for head-fixation of the animal, was attached to the window glass coverslip.

**B.** Tile scan of thalamic injection site with projections reaching up to the cortex (wide field epifluorescence, scale bar 1 mm).

**C.** GFP-labelled mitochondria in thalamic projections imaged in upper layers of the cortex (two-photon, scale bar 25 μm).

**D.** Immunofluorescence staining of GFP-labelled mitochondria in neurons of the MD thalamus co-stained with the outer mitochondrial membrane marker TOM20 (single confocal plane, scale bar 5 μm).

**E.** Mean velocity under basal conditions was 0.30 ± 0.20 μm/s (n = 406 mitochondria from 4 mice). Data displayed as Box-Whisker Plot; mean indicated as +, whiskers show min-max. Reference dots: red (Kiryu-Seo et al., 2010) (*in vitro*); light blue (Takihara et al., 2015) (*in vivo*).

**F.** Time course of mitochondrial velocity under basal conditions (black) and after ethanol injection (green). Mitochondrial velocity was unaffected by acute ethanol intoxication (p = 0.4601, two-way repeated measures ANOVA, n = 8 focal planes from 4 mice; mean ± SEM).

**G.** Histogram of mitochondrial velocity with fit (R^2^ = 0.9578) showing the sum of two Gaussian distributed populations of mitochondria transported at different mean speeds.

**H.** Mean DCV velocity under basal conditions was 0.48 ± 0.2 μm/s (n = 4657 DCVs from 3 mice). Data displayed as Box-Whisker Plot; mean indicated as +, whiskers show min-max. Reference dots: dark blue (de Wit et al., 2006) (*in vitro*); grey (Kwinter et al., 2009) (*in vitro*); brown (Knabbe et al., 2018) (*in vivo*).

**I.** Time course of DCV velocity under basal conditions (black) and after ethanol injection (green). DCV velocity increased after acute ethanol intoxication as well as in the saline condition; the increase was not significantly different between saline and ethanol injected mice (n = 3, mean ± SEM).

**J.** Kymographs of the same axonal stretch before and 210 min post-ethanol injection. The number of transported DCVs remains stable.

**K.** Representative images of mitochondria (orange arrow heads) that delocalized/disappeared from the imaged region 4 h after ethanol application (*in vivo*, scale bar 5 μm).

**Supplementary Figure 4: Acute ethanol intoxication does not affect mitochondrial structure, but mitochondrial occupation of presynaptic boutons**

**A.** *In vivo* occupied and unoccupied presynaptic boutons (red), mitochondria depicted in grey (scale bar 5 μm)

**B.** 80.5 ± 9.1 % of presynaptic boutons (n = 80 axons from 4 mice) were occupied by mitochondria in thalamic long-projection neurons. 4 h after ethanol intoxication occupancy of presynaptic boutons was significantly reduced (P=0.0163). After 24 h occupancy returned to baseline level (P<0.0001) (Two-way repeated measures ANOVA with Sidak’s multiple comparisons test, F (1,78) = 0.1315).

**C.** Diameter of stable, lost and formed occupied (occ.) and unoccupied (unocc.) presynaptic boutons (n > 50 per group). Occupied stable presynaptic boutons were significantly larger than other boutons. Occupied boutons had on average a larger diameter (1.26 μm) than unoccupied boutons (0.89 μm) and also reached higher maximal diameters (3.03 μm) compared to unoccupied synapses (2.372 μm) (P<0.0001, Kruskal-Wallis test, Dunn’s multiple comparison test for differences between occupied boutons and all other boutons, differences between the rest are not significant).

**D.** Mean mitochondria length was 1.52 ± 0.73 μm (n = 1343 mitochondria from 4 mice). Data displayed as Box-Whisker Plot; mean indicated as +, whiskers show 1-99 percentile.

**E**. Mitochondrial length (n > 600 mitochondria per time point from 4 mice, mean ± SEM) was unaffected by acute ethanol intoxication (p = 0.17, two-way repeated measures ANOVA).

**Supplementary Figure 5: Dynamics of mitochondrial occupation of presynaptic boutons.**

**A-C.** Longitudinal *in vivo* imaging of boutons (marked in red by SyPhy-mCherry) and mitochondria (white, mitoGFP) revealed the dynamics of bouton turnover with or without mitochondria.

**D.** Loss of occupied presynaptic boutons was unaffected by acute ethanol intoxication when comparing the saline to the ethanol condition (P=0.822), however, a significant increase after 24 h compared to 4 h could be detected in the ethanol condition (P=0.0072), but not in the saline condition (Two-way repeated measures ANOVA with Sidak’s multiple comparisons test, F(1,78) = 0.05087).

**E.** Formation of occupied presynaptic boutons was unaffected by ethanol intoxication (p = 0.95, two-way repeated measures ANOVA, F (1,78) = 0.004359).

**F.** Formation of unoccupied presynaptic boutons was unaffected by ethanol intoxication (p = 0.1433, two-way repeated measures ANOVA, F (1,78) = 2,186).

**Supplementary Figure 6: Time course of behavioral inhibition after a single dose of ethanol**

After achieving a stable performance for the Go/NoGo task in two training sessions, mice were retested for different variables in experimental sessions at 4-6 h, 24 h, and 48 h following injection with 3.5 g/kg ethanol. (**a**) Correct go: number of right responses to go (4-6 h: *t*_(5)_=4.1, P<0.01; 24 h: *t*_(5)_=2.3; P=0.07; 48 h: *t*_(5)_=1.7; P=0.1). (**b**) False alarm: number of going when they should not go (4-6 h: *t*_(5)_=2, P=0.1; 24 h: *t*_(5)_=2.1, P=0.148; 48 h: *t*_(5)_=2.6, P<0.05). (**c**) Efficiency: total number of reinforcers earned/total number of lever presses (4-6 h: *t*(5)=3.2, P<0.05; 24 h: *t*_(5)_=2.7, P<0.05; 48 h: *t*_(5)_=1.6, P=0.2) (**d**) Performance rate: correct go/active lever pressings (4-6 h: *t*_(5)_=4.9, P<0.01; 24 h: *t*_(5)_=3.8, P<0.05; 48 h: *t*_(5)_=2.2, P=0.1). All values are given as mean ± SEM, and statistical significance was set at P < 0.05.

**Supplementary Table 2: Odours/odorants preferences of trained adult flies.**

The RNAi transgenes used are *UAS-milton-RNAi^GD8116^, UAS-milton-RNAi^TRiPJF03022^* and *UAS-dmiro-RNAi^TRiPJF02775^*. The errors are s.e.m.

* The values differ significantly from random choice as determined using One-sample sign test.

** None of the values differ from random choice as determined using One-sample sign test and they did not differ between the controls and experimental groups as determined using ANOVA *post-hoc* Tukey-Kramer HSD.

### Supplementary Information

#### Protein candidates identified by mass spectrometry

Overall, 72 significantly changing proteins were detected of which 32 proteins were affected in both young and old mice (**Table 1**). These changes in abundance may be a consequence of distinct cellular mechanisms including changes in protein turnover dynamics, newly synthesized proteins or protein trafficking in and out of the synapse. Note that we did not intend to characterize the underlying causes for the abundance change but rather aimed to identify molecular candidates that mediate some of the complex alcohol-related effects in the brain.

Of the 72 proteins, 39 changed significantly in 2 independent experiments (in young or old animals; p<0.0025), 24 proteins changed significantly in 3 independent experiments (p<0.000125), and 9 proteins changed significantly in all 4 independent experiments (p<6.25*10^−6^). This high reproducibility of the *ex vivo* approach therefore suggests that it is suitable for identification of proteins that change their synaptic abundance upon ethanol exposure.

Unlike chronic ethanol consumption, acute ethanol exposure predominantly manifests itself in complex behavioral responses that depend on alterations of neuronal transmission. We detected an increase in Clathrin light chains A and B, both are key components of the recycling synaptic vesicle, and an increase in the kinase SAD1 which was found to regulate synaptic transmitter release (Inoue et al., 2006). In addition, ethanol led to an increase of two vacuolar ATP synthase subunits involved in acidification of synaptic vesicles which in turn aids loading the vesicles with neurotransmitter via the proton gradient. Ethanol also induced an increase of the Freud-1 repressor at the synapse, which was found to inversely regulate dopamine D2 receptor (Rogaeva et al., 2007) and serotonin 5-HT1A receptor levels (Ou et al., 2000). The dopamine D2 receptor has been linked to genetic predisposition for alcoholism while the serotonin receptor is being implicated in mood regulation and psychiatric disorders. Several proteins linked to antidepressant treatment were also affected including cAMP-specific 3’,5’-cyclic phosphodiesterase 4D (PDE4D), a key target of the antidepressant Rolipram (Zhang et al., 2002). In addition, we discovered an ethanol-dependent elevation in monoamine oxidase A, deficiencies of which are associated with heightened aggressiveness (Alia-Klein et al., 2008). A pronounced change (2.19) was seen with MAP6 which elicits various cognitive impairments in null mutant mice (Andrieux et al., 2002).

Acute consumption of ethanol has long been known to diminish memory formation both in humans as well as animal models and pharmacologically relevant concentrations (50-100 mM) of ethanol inhibit long-term potentiation (LTP) (White et al., 2000). The underlying molecular and cellular mechanisms of this impairment are not well understood and only a few proteins implicated in the inhibition of LTP have been described (Wang et al., 2007). We identified six additional synaptic proteins that could potentially play a role in the acute ethanol-mediated impairment of LTP. These include Tenascin-R (Bukalo et al., 2007), GAP-43 (Hulo et al., 2002), MAP6 (Andrieux et al., 2002), SNAP-25 (Hou et al., 2006), Phosphacan (Takeda-Uchimura et al., 2015), and Ankyrin-G (Smith et al., 2014). In addition, a reproducible, but not statistically significant decrease of a key plasticity protein, the NMDA receptor 2B (0.91) was also detected at P30.

Ethanol is also known to elevate GABA-mediated synaptic inhibition but the molecular basis for this is not fully understood (Weiner and Valenzuela, 2006). We found a decrease of GABA transaminase, which is necessary for GABA catabolism, and a decrease of GABA transporter 4 levels (**Supplementary Fig. 2B**), a protein involved in recycling released GABA transmitter from the synapse. The combination of both could at least partly explain the observed inhibitory action of ethanol.

We detected an increase of C-terminal binding protein 1, which positively affects expression of the multi-drug resistance protein (Jin et al., 2007), a finding that potentially has important medical implications. Similarly, expression of beta tubulin III, a pan-neuronal marker for differentiated neurons, was increased in all four experiments while overexpression of beta tubulin III in tumors has been associated with low chemotherapy response rates when using tubulin-binding agents (Ferrandina et al., 2006). Also to our surprise, the moderate ethanol exposure was sufficient to affect synaptic abundance of proteins connected to apoptosis (Apoptosis inducing factor (AIF) (Krantic et al., 2007), Adenylyl cyclase-associated protein 1 (CAP1) (Wang et al., 2008), asparagine synthetase (Cui et al., 2007), thioredoxin reductase 2 (Nonn et al., 2003)). Ethanol has often been linked to apoptosis in the developing brain, especially in case of the fetal alcohol syndrome (FAS), but little is known about the apoptotic effects of acute exposure in the adult brain and even less is known about the underlying mechanisms. Our data highlight potential molecular pathways that could lead to programmed cell death in mature, differentiated neurons.

We detected a total of 19 ATP synthase proteins in the hippocampal synaptic proteome and 13 of these were significantly changing upon ethanol treatment. For the mitochondrial ATP synthase, we found an almost stoichiometric increase 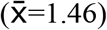 with a remarkably low variance (STD=0.05) for all 8 identified subunits suggesting that this ATP synthase complex is affected in its entirety rather than each protein changing independently. As expected, the change in abundance was inverse for certain proteins after acute versus chronic ethanol exposure (e.g. malate dehydrogenase).

Overall, there is a tremendous disconnection between our detailed understanding about the various forms of neuronal plasticity and our lack of knowledge about the changes in molecular compositions that mediate it. On a molecular level, we show here that high accuracy detection of minute, stimulus-dependent changes of synaptic protein abundance is now possible, including changes related to synaptic plasticity (SBC, unpublished results). Prior to this study, the protein correlate of synaptic plasticity may have been imagined anywhere from a handful of ‘Master’ proteins to a large fraction of proteins. Our data provide a first benchmark for the number of stimulus-dependent protein abundance changes with 3.5% of the total synaptic proteome.

## Methods

### Ethical approval

Animal experiments were carried out in accordance with the EU and German guidelines for the care and use of laboratory animals and were approved by the Governmental Supervisory Panel on Animal Experiments of Baden Wuerttemberg in Karlsruhe (Aktenzeichen 35-9185.81/G-166/15 and 35-9185.81/G-24/15). Priority was given to avoid or minimize animal suffering, while promoting animal welfare and reducing the number of animals using the 3R principle, for example through longitudinal experiments. Mice were housed singly in an individually-ventilated cage system (ZOONLAB) at a 12 h dark/light cycle. The animal room was tempered to 21-23 °C with a relative humidity of 55 %. Mice had access to water and chow *ad libitum*, except for experimental sessions. Behavioral tests were performed in the dark cycle. Mice used for behavioral tests were handled daily before starting the experiments and were habituated to the behavioral testing environments. Mice that were exposed to ethanol were between ages 2-7 months.

### SILAC mice

The generation of fully labeled SILAC mice was described previously (Kruger et al., 2008). Briefly, a SILAC-diet was prepared by adding ^13^C_6_-lysine to a customized lysine-free mouse diet (Harlan) to a final concentration of 1% according to standard mouse nutritional requirements. After feeding two generations of female C57BL/6N mice with the ‘heavy’ diet, complete *in vivo* labeling (>97 %) was achieved in the F2 offspring which contained virtually no unlabeled peptides (Kruger et al., 2008). Only females were maintained from each litter and fed with the ‘heavy’ diet so that the offspring also fully incorporated the ^13^C_6_-lysine. One animal was available for the PND30 experiments, one for PND210. The experiments were performed with animals from the F2 or F3 generation and labeled animals showed normal breeding behavior and motor activity.

### Hippocampal slice preparations and synaptic protein extraction

Acute hippocampal slices were prepared according to standard procedures (Lein et al., 2011). Briefly, mice were anesthetized, rapidly perfused with ice-cold saline, and the brains quickly transferred to ice-cold carbogenated (95% O_2_, 5% CO_2_) artificial cerebral spinal fluid (ACSF in mM: 124 NaCl, 5 KCl, 1.25 Na_2_PO_4_, 26 NaCO_3_, 2.5 CaCl_2_, 2 MgSO_4_, 10 Glucose). Hippocampal slices (400 μm) were allowed to recover in ACSF at room temperature for 1 h and were then transferred to ACSF with or without 50 mM ethanol (29-31° C; 4 h). Slices were combined in a cross-over fashion and homogenized in 0.32 M sucrose, 5 mM HEPES pH7.4, protease inhibitors (Roche tablets, EDTA free). The homogenate was centrifuged (10 min/1000 g) and the supernatant re-centrifuged (20 min/ 12000 g). The pellet was resuspended in homogenization buffer, layered onto a sucrose gradient (0.85 M, 1.0 M, 1.25 M; all solutions contained 5 mM HEPES pH7.4, protease inhibitor), and centrifuged (2 h/ 82500 g). Synaptosomes were collected at the 1.0/1.25 M interface and adjusted to a final concentration of 20 mM Tris pH6 and 1 % Triton-X (Phillips et al., 2001). Synaptosomes were extracted for 30 min and subsequently centrifuged (15 min/ 15000 g). Pellets were re-extracted in 20 mM Tris pH6/ 1 % Triton-X and re-centrifuged (15 min/ 15000 g). The resulting pellet was dissolved in Laemmli buffer and directly loaded on a 1D SDS gel.

### In-gel digest and peptide extraction

In-gel digest was done according to standard procedures (Kruger et al., 2008). Synaptic proteins were separated on a 4-12 % NuPage Novex Bis-Tris gel (Invitrogen), the gel stained using the Colloidal Blue Staining Kit (Invitrogen) and cut into 10 slices. The in-gel digest was performed with an overnight digestion at 37° C using 12.5 nmol endopeptidase LysC. Each sample was loaded on C_18_ StageTips, washed, and eluted.

### LC-MS/MS

All LC-MS/MS experiments were performed by standard procedures (Kruger et al., 2008). Briefly, peptides were separated using an Agilent 1100 or 1200 nanoflow LC-System. The HPLC system was coupled to an LTQ-Orbitrap mass spectrometer (ThermoFisher Scientific). Survey full scan MS spectra (m/z 300-2000) were acquired in the orbitrap analyzer. The five most intense ions from the survey scan were sequenced by collision induced dissociation in the LTQ. Data were acquired using Xcalibur software.

### Bioinformatic analysis

Mass spectra were analyzed using the in house developed software MaxQuant (version 1.0.12.5) (Cox, 2008). The data were searched using Mascot (version 2.2.04, Matrix Science) against the Mouse International Protein Index protein sequence database (IPI, version 3.37) supplemented with frequently observed contaminants and concatenated with reversed copies of all sequences (2 x 51,467 entries). Initial maximum allowed mass deviation was set to 5 ppm for monoisotopic precursor ions and 0.5 Da for MS/MS peaks. The required false positive rate was set to 1% at the peptide level, the required false discovery rate was set to 1% at the protein level and the minimum required peptide length to 6 amino acids. In addition to the protein false discovery rate threshold, for protein identification we required at least two unique sequenced peptides, while for protein quantitation, at least two independently sequenced and quantified SILAC pairs, one ‘forward’ and one ‘reverse’ in the respective cross-over experiment were necessary. We allowed three missed cleavages and the enzyme specificity was set to Lys-C. We included carbamidomethylcysteine as a fixed modification, and *N*-acetylation and methionine oxidation as variable modifications. Contaminants such as keratins were discarded. The analysis of protein significance is described in detail elsewhere (Cox, 2008). Briefly, the significance of protein ratios was computed taking into account the summed peptide intensity (‘Significance B’ of the MaxQuant software). To assess if the ethanol-dependent change of a given protein was significant, the complementary error function was used to produce a probability measure (‘p’) of the fold change. Statistically, only proteins that displayed a significant change in both, forward and reverse ratios (P < 0.05; P × P <0.0025) were included. Furthermore, with a minor extension of our statistical requirements (P < 0.075; P × P < 0.0025) 17 additional proteins that merit consideration based on their biological roles also became statistically significant, including GAP-43 and Fyn kinase, both reported to be affected by ethanol (Kim et al., 2006; Miyakawa et al., 1997). Those were included in the list of the 72 proteins that changed significantly upon ethanol exposure.

### *In vivo* ethanol exposure

For all experiments with living mice, we based our research on an extensive body of publications (Eisenhardt et al., 2015; Haseba et al., 2012) showing that mice have a high ethanol metabolism, which reduces blood and breath alcohol concentrations to negligible levels within 3-4 h, including initial ethanol doses of > 3 g/kg (Ginsburg et al., 2008). Thus, the assumption is that any ethanol-dependent change seen 4 h after injection is reflective of a lasting effect of the initial ethanol dose.

### Immunofluorescence of mitochondria and dense core vesicles

At different time points after the mice had received a single dose of ethanol (3.5 mg/kg body weight), mice were transcardially perfused with 4 % PFA fixative and subsequently the brains were carefully removed from the skull. For immunofluorescence stainings, 80 μm coronal sections of the S1/M1 cortex were cut with a microtome (Leica VT100S, Leica). Brain sections were first blocked in PBS containing 5 % natural goat serum (NGS) and 1 % Tritum-X100 for 2 h, before incubating the slices in PBS containing 1 % NGS, 0.2 % Triton-X100 and the primary antibody over night at 4 °C. Slices were washed in PBS containing 2 % NGS. Secondary antibodies were incubated for 2-3 h at room temperature. Primary antibodies: α-Translocase of the outer mitochondrial membrane (TOM20, 1:500; Santa Cruz), α-Synaptophysin 1 (SyPhy1, 1:500; Synaptic Systems), α-Chromogranin A (Chr-A, 1:500; Synaptic Systems). Alexa Fluor antibodies (1:500; Invitrogen) were used as secondary antibodies. Slices were mounted with Mowiol (SigmaAldrich) and imaged on a Leica DM6000 wide-field epifluorescence light microscope with 10x/0.4 (magnification/NA) or 63x/1.30 glycerol-immersion objectives or on a Leica SP8 scanning confocal microscope using a 63x/1.40 oil-immersion objective.

### Immunofluorescence for detection of synaptic protein dynamics *in vivo*

300 nm cryosections were washed in PBS 3 x 10 minutes. The cryosections were blocked in 0.5 % inactivated fetal calf serum (FCS). The primary antibody was added at the desired concentration in 0.5 % inactivated FCS. Primary antibodies were incubated for 45 minutes at 4 °C followed by three washing steps in PBS for 10 minutes. The secondary antibody was added at the desired concentration in 0.5 % inactivated FCS for 1 hour at 4 °C. CS were washed in PBS 3 x 10 minutes. Subsequently, CS were mounted with Polymount. For the immunofluorescence co–localization analyses, cryosections were stained for β–Actin (Alexa488), Synapsin (Alexa647), and either MAP6, Ankyrin-G, or PCCA (Alexa594). The resulting single 8–bit Tiff images were thresholded individually. Both, the Synapsin and the Actin signals were used to identify putative synaptic structures within the image, but the Synapsin signal was ultimately used for normalization of MAP6, Ankyrin-G, or PCCA signals. To count the overlapping structures between the Synapsin and the MAP6/PCCA/Ankyrin-G staining those two channels were subtracted from each other. The resulting image now contained only binary ROIs of overlapping Synapsin and MAP6/PCCA Ankyrin-G regions. Those ROIs were plotted on the unprocessed images and the fluorescence values in these regions was measured with ImageJ. Expression ratios were then calculated from the intensity of the protein of interest divided by the Synapsin intensity in all overlapping ROIs found in each image. Synapsin (1:100, Synaptic Systems, Company ID: 106 004), β-Actin (1:200, Santa Cruz, Company ID: sc-28561), MAP6 (1:200, Santa Cruz, Company ID: sc-53513), PCCA (1:200, Santa Cruz, Company ID: sc-374341) and Ankyrin G (1:200, Santa Cruz, Company ID: sc-28561)

### Immunofluorescence for AIS length analysis

Mice were transcardially exsanguinated under deep anesthesia with Ketamine (120 mg/kg BW) / Xylazine (16 mg/kg BW) with 0.9% saline followed by ice-cold 2 % paraformaldehyde (Roth; in 0.1 M PBS, pH 7.4) and brains removed and cryoprotected in 30% sucrose in PBS at 4° overnight. Samples were then trimmed to a block containing S1, embedded in Tissue Tek^®^ (Sakura Finetek) and frozen in liquid nitrogen-cooled isopenthane (Roth). Cryosections (30 μm) were cut using a cryotome (Microm HM 550, Thermo Fischer) and immediately processed for free-floating immunofluorescent staining. After a brief rinse in PBS, slices were blocked in fish skin gelatin blocking buffer (1% BSA, 0.2% fish skin gelatine, 0.1% Triton in 0.1 M PBS) for at least 90 min and subsequently incubated in primary antibody diluted in blocking buffer (4°C overnight) using antibodies against the AIS scaffolding protein βIV-spectrin (self-made rabbit polyclonal antibody against amino acids 2237-2256 of human ßIV-spectrin; 1:500) (Schluter et al., 2017) and the neuronal nuclei marker NeuN (mouse monoclonal, Millipore; 1:500). Omission of the primary antibodies did not yield specific immunolabeling. After primary antibody incubation, slices were rinsed 4 x 10 min in PBS at room temperature and then incubated in secondary antibodies (goat anti mouse Alexa Fluor 568, 1:1000 and goat anti rabbit Alexa Fluor 488, 1:1000; Molecular Probes, Thermo Fisher) for 120 min in the dark. Slices were then rinsed 4 x 15 min in PBS using the nuclear counterstain TO-PRO-3 Iodide (1:2000, Thermo Fisher). For preservation of immunofluorescence, slices were mounted in a mounting medium with anti-fading effect (Roti-Mount FluorCare, Carl Roth) and stored at 4°C.

### Image acquisition and analysis of AIS length

Confocal analysis was carried out on a C1 Nikon confocal microscope with a 60 x objective (oil immersion, numerical aperture of 1.4; Nikon Instruments). To increase the number of infocus immunoreactive AIS, maximum intensity projections were saved and processed. Thickness of single optical sections was 0.5 μm in stacks of 10 - 20 μm total depth. Confocal x-y-resolution was kept at 0.21 μm per pixel. Images for qualitative analysis were evaluated and enhanced for contrast in Photoshop (Adobe Systems, USA). AIS length was measured using the self-written software AISuite (Ernst et al., 2018; Hofflin et al., 2017), which is available online (github.com/jhnnsrs/aisuite2). This tool extends the well-established and widely used method of defining AIS start and end points as points where a predefined fluorescence threshold (relative to the maximum fluorescence intensity along a line drawn over an individual AIS) is surpassed (Grubb and Burrone, 2010). Using AISuite, the threshold was adjusted depending on the individual staining quality in each section and ranged from 10 - 30% of maximum fluorescence intensity.

### *In vitro* overexpression of candidate proteins

Rat hippocampal cultures were prepared at embryonal day 18.5 as described (Kaech and Banker, 2006). Plasmids of GFP or GFP fusions with the 190 kD AnkG version (Smith et al., 2014) or full length MAP6-N (Peris et al., 2018) were co-transfected with a control plasmid expressing mOrange2 under the neuronal Synapsin promoter. Transfections were performed after 8 days *in vitro* (DIV8) with Lipofectamine2000 according to the manufacturer’s protocol, except using 1 μg instead of 0.5 μg of plasmid DNA together with 1 μl of Lipofectamine per well in 24-well plates. At DIV13, half of the cultures were exposed to 50 mM ethanol for 4 h. Following fixation with 4% PFA for 10 min, a blinded observer imaged hippocampal neurons with dual GFP and mOrange2 expression with a confocal microscope and manually counted the spines using the red mOrange2 channel. Image analyses were done with Fiji.

### Spine analysis following ethanol exposure *in vivo*

Thy1-GFP mice (Feng et al., 2000) were co-injected with pyrazole (Sigma) in saline at 1 mmole/per kg mouse (Eisenhardt et al., 2015) and ethanol at 3.5 g/kg (from a 20% solution) or a corresponding volume of saline for control mice. Ethanol treated mice were additionally kept in an ethanol vapor chamber (La Jolla Alcohol Research, California, USA) (Hirth et al., 2016). A peristaltic pump delivered 98 % alcohol at a flow rate of 0.44 ml/min to a heated roundbottom flask so that the alcohol quickly vaporized. Pressure controlled airflow to the flask carried the alcohol vapor to the individual chambers which were connected to a vacuum for constant air exchange. The airflow was adjusted to about 6 l/min to maintain the ethanol concentration of 10-15 mg/l air in the chamber. After 4-6 hours, mice were perfused with 4 % PFA and the brains sliced. Dendritic spines on cortical dendritic stretches of GFP-positive neurons were imaged with a confocal microscope and manually quantified by a blinded observer.

### *In vivo* two-photon imaging of synaptic turnover, mitochondria and DCVs in mice

#### Genetic labelling of mitochondria, DCVs and presynaptic boutons

In house produced (Schwenger and Kuner, 2010) recombinant adeno-associated viruses (rAAVs) of the chimeric 1/2 serotype were used in order to label axonal organelles and presynaptic terminals in neurons *in vivo*. Mitochondria were labelled with GFP fused to the first 35 amino acids of the mitochondria-targeting sequence of cytochrome *c* oxidase subunit VIII leading to enrichment of GFP in the mitochondrial matrix (Plasmid was kindly provided by Dr. Ken Nakamura, see Berthet et al. 2014). Mitochondria-targeted GFP (mitoGFP) was expressed under the control of the EF-1 alpha promoter. As the construct contained a Cre-inducible double-floxed inverse open reading frame (DIO), Cre-recombinase was necessary for expression of the insert (EF-1α-DIO-mitoGFP). rAAVs encoding for Cre-recombinase under the control of the synapsin promoter (Syn-Cre) were always co-injected leading to a restricted expression of mitoGFP in neurons. For the labelling of dense-core vesicles (DCV), the DCV cargo protein neuropeptide-Y (NPY) fused to the fluorescent protein Venus was used under the control of the Synapsin promoter. The plasmid for Syn-NPY-Venus was kindly provided by Dr. Matthijs Verhage. DCV labelling was combined with axonal labelling by expressing mCherry under the CAG promoter (Syn-mCherry). Labelling of presynaptic boutons was achieved by expressing synaptophysin (SyPhy) fused to mCherry under the control of the CAG promoter (CAG-SyPhy-mCherry).

### Craniectomy, viral injection, and chronic cranial window implantation

Protocol was adapted from (Holtmaat et al., 2009) as well as (Boffi et al., 2018). Mice were deeply anaesthetized by intraperitoneally (i.p.) injecting a mixture of 0.48 μl fentanyl (1 mg/ml; Janssen), 1.91 μl midazolam (5 mg/ml; Hameln) and 0.74 μl medetomidin (1 mg/ml; Pfizer) per gram body weight each and mounted in a stereotaxic apparatus (EM70G, David Kopf Instruments). Mice were kept on 35°C by a feedback-controlled heating pad throughout the surgery. The eyes were covered with eye ointment (Bepanthen with 5% Dexpanthetol, Bayer Vital GmbH) to prevent drying-out. Standard aseptic procedures were followed during all surgeries. The local anesthetic Xylocain (1%, AstraZeneca) was injected subcutaneously (s.c.) at the site of surgery and the cranium was exposed by removing a flap of the scalp, approximately 1 cm2, as well as the subjacent gelatinous periosteum. Levelling of the mouse skull to a coordinate system centered at bregma was performed by eye using a monocle and a digital indication (both David Kopf Instruments). To target the somatosensory cortex of the right hemisphere in Thy1-GFP mice (Feng et al., 2000) following coordinates where used relative to the bregma (in mm): −2.5 AP, 2 ML. Since Thy1-GFP mice endogenously express GFP in neurons, those mice were only implanted with cranial windows, without viral injections. In contrast, in wt mice a small hole was drilled into the skull with a dental drill (drill: EXL-40, Osada Inc.; boring head: 1104005, Komet Dental) and the right-hemisphere medio-dorsal nucleus of the thalamus (MD thalamus) was targeted using the coordinates 0.82 AP, 1.13 ML and 3.28 DV (Paxinos and Franklin 2013) mm relative to bregma. Approximately 800 nl of a mixture of rAAVs (1:1:1 mitoGFP, Cre, SyPhy-mCherry OR 1:1 NPY-venus, mCherry) was slowly injected. For cranial window implantation, a circular part of the skull with a diameter of 6.5 −7 mm (centered above the injection site in wt mice) was removed by drilling a deep groove into the outline of the circle. The dura mater was carefully removed with very fine forceps (straight Dumont #5, Fine Science Tools Inc.). A glass coverslip (6 mm diameter, #0) was placed inside the opening and was, together with a 3D-printed round plastic holder, cemented to the scull will dental acrylic cement (glue: Cyano, Hager Werken; powder: Paladur, Heraeus). Mice received i.p. a mixture of 1.86 μl naloxon (0.4 mg/ml; Inresa), 0.31 μl flumazenil (0.1 mg/ml; Fresenius Kabi), 0.31 μl antipamezole (5 mg/ml; Pfizer) in 3.72 μl saline (0.9 %; Braun) each per gram body weight to antagonize the anesthesia. For pain treatment, mice received an s.c. injection of 150 μl Carprofen (50 mg/ml; Bayer Vital GmbH) diluted in saline every 12 h for the next two days. Mice rested for 5-8 weeks to ensure high rAAV expression and an eased inflammatory glial reaction (Holtmaat et al., 2009).

### In vivo two photon microscopy of anesthetized mice and ethanol injection

Two photon imaging (Denk et al., 1994) was conducted as described in (Knabbe et al., 2018) using an upright TriM Scope II microscope (LaVision BioTec GmbH) equipped with a pulsed Ti:Sapphire laser (Chameleon Ultra II; Coherent). A wavelength of 960 nm was used to excite Thy1-GFP OR mitoGFP OR simultaneously NPY-venus and cytosolic mCherry. Imaging of mitoGFP with 960 nm and SyPhy-mCherry with 1050 nm was serially done. The emitted signals were separated using a 560 nm dichroic mirror (Chroma, Bellows Falls) and appropriate filter sets (GFP, venus: 535/70 nm emission filter (Chroma, Bellows Falls); mCherry: 645/75 nm emission filter (Semrock, Rochester)). Imaging was done using a 25x water immersion objective (1.1 NA, Nikon) and the emitted fluorescence was collected with photo multiplier tubes (PMTs, H7422-40-LV 5M; Hamamatsu). Mice were anesthetized in an acrylic box with 5 % isoflurane (Henry Schein) in medical O_2_ (Air Liquid Medical GmbH) using a vaporizer (Vapor 2000, Dräger). The animal was head-fixed into a custom-built holder screwed onto the intravital microscope stage. Vessels were used for orientation and relocating to the same imaging site during every imaging session. During the imaging session, isoflurane was maintained at 0.8 – 1.5%. Breathing rate was under constant monitoring using an infra-red camera (ELP 1080P 2.0 Megapixel USB camera, Ailipu Technology Co.) and an infra-red LED light source (48 LED Illuminator, Sonline). Body temperature of the animal was kept at 37 °C by a heating pad (ExoTerra, HAGEN Deutschland GmbH).

For imaging spines in Thy1-GFP mice, imaging sites were chosen in such a way that enough dendritic stretches in S1/M1 layer II/III were visible und feasible for post hoc spine counting. Since Thy1-GFP mice show very dense labeling in this layer we aimed for sparsely labeled regions. 3D-stacks covering 436 x 436 x 50 μm, at 2731 x 2730 pixel resolution (0.16 μm/pixel resolution) and 1 μm z-steps were acquired starting from the pial surface. ImSpector (LaVision BioTec GmbH) was used for microscope control and image acquisition. Mice were imaged on three consecutive days (day 1 = pre-ethanol, day 2 = ethanol, day 3 = post-ethanol). Each day included 4 time points: 0, 2, 4 and 8 h. Mice were kept in their cages between the imaging sessions to minimize stress. On day 2, mice received an i.p. injection of ethanol (3.5 g/ kg) at time point 0 h, and a second ethanol injection at 2 h. On the pre-ethanol and post-ethanol days mice received a saline injection.

For imaging mitochondria, DCVs or pre-synaptic boutons, layer 1 of motor cortex, 50-100 μm away from the pial surface was imaged. When imaging mitochondria, frames were taken from an area covering 436 x 436 μm, at 1024 x 1024 pixel resolution (0.426 μm/pixel resolution) with a frame rate of 0.61 Hz using a galvanometer scanner. During acquisition of mitochondrial transport the focal plane was rapidly alternated using a piezo-motor (LaVision BioTec GmbH) allowing the acquisition of three focal planes at each time point resulting in an effective frame rate of 0.2 Hz per image plane. When imaging DCVs, frames were taken from an area covering 200 x 200 μm, at 1071 x 1071 pixel resolution (0.19 μm/pixel resolution) with a frame rate of 0.94 Hz using the galvanometer scanner. Each acquired time-lapse of mitochondrial transport was of 10 min duration and 5 min of DCV transport. Baseline time-lapses were recorded at three time points before i.p. injection of 500 μl of saline or 20 % ethanol in water. Time-lapses up to 4 h post-injection were acquired. Three additional time-lapses were acquired 24 h post-ethanol injection. 3D-stacks covering 436 x 436 x 50 μm, at 2731 x 2730 pixel resolution (0.16 μm/pixel resolution) and 1 μm z-steps were acquired pre, 4 h and 24 h post saline and ethanol injection in order to dissolve structural changes of mitochondria and presynaptic terminals.

### Post hoc analysis of synaptic turnover

3D-stacks of all time points were registered by manual selection of corresponding points and calculation of the resulting transformation in Matlab using a customized script. For analyzing spine turnover, 5 dendritic stretches of approx. 50 μm length were chosen randomly for analysis in each mouse. For analyzing the structural plasticity of mitochondria and presynaptic terminals axons clearly visible in all time points (pre, 4 h and 24 h post) were chosen. Image processing was done in the open-source image analysis software Fiji. Dendrites and axons were analyzed in 3D and searched for spines/boutons manually. 3D stacks covering 436 x 436 x 500 μm ROIs where cropped in x, y spanning the dendritic stretches of interest. In z, the image was cropped to 25 μm above and below the center of the dendrite. Spine labeling, counting and calculation of the spine turnover ratio (TOR) was performed according to previously published procedures (Holtmaat et al., 2005): Spines were considered lost when they retracted into the dendrite (length < 5 pixels) or considered gained when they formed clear protrusions from the dendrite (length > 5 pixels). TOR was calculated for each day individually as (N_gained_ + N_lost_)/(2xN_total_) where N_gained_ is number of spines newly formed between two consecutive time points (eg. Time point 0 h and 2 h, 2 h and 4 h, 4 h and 8 h), whereas N_lost_ is the number of spines lost between the consecutive time points. N_total_ represents the absolute number of spines counted in one time point. This results in 3 TOR values for each day, corresponding to the spine turnover from 0 to 2 h, 2 to 4 h and 4 to 8 h, respectively. Mean TOR for each dendritic stretch was calculated per day taking the mean out of the 3 TOR values for each of the 3 imaging days. All 3 days where compared statistically with each other. Spine density was calculated as spines/μm. The mean spine densities of the 4 time points (0, 2, 4 and 8 hours after injection of ethanol) were calculated for each dendritic stretch and statistically compared using a repeated measures ANOVA and a post hoc Bonferroni’s test for multiple comparisons. For boutons, signals 1.5 times higher than the background of the axon were counted as mitochondria or presynaptic terminals, respectively. Turnover rate of boutons was calculated by comparing numbers of lost/gained boutons 4 h and 24 h post ethanol/saline to existing boutons in the pre ethanol/saline time point.

### Analysis of mitochondrial and DCV transport

For image processing and analysis, the open-source image analysis software Fiji was used. Time-lapse images were registered in *xy* using the Fiji plugin “moco” (Dubbs et al., 2016). Average intensity projections of time-lapses were used to show stationary cell organelles as well as the outlines of axons, while maximum intensity projection was used to identify axons displaying active cell organelle trafficking. For mitochondrial transport, only active axonal stretches (ROIs) for analysis were selected. A total of 15 ROIs was analyzed per focal plane. Kymographs were generated with the Multi Kymograph tool (line width = 3) in Fiji. Kymographs revealed the number of stationary and mobile mitochondria, from which a mobility ratio was calculated. Kymographs were used to calculate the mean velocity of a mitochondrial track by using a Fiji macro written by Alessandro Moro. DCV transport was automatically tracked (see next section) and analyzed using self-written routines in Matlab (MathWorks). For mobility, DCV tracks were defined as mobile, if the total movement distance was above or equal to 5 μm and stable if the movement distance was below 5 μm. Mobility was calculated as mobile-fraction / (stable fraction + mobile fraction). Mean velocity was calculated on the mobile fraction only.

### Automatic tracking of dense core vesicles

To quantify the mobility of vesicles in the mouse brain data, an automatic tracking approach was used (Jaiswal et al., 2015). In each image frame of a video, vesicles were detected by applying the spot-enhancing filter (Dubbs et al., 2016). Two different size thresholds yielding two sets of detections at each time point were used. Detected spots of small and large size were used for tracking, while for initialization of trajectories only large spots were used. A Kalman filter (Kalman, 1960) was employed to determine predictions about the state of vesicles using a two motion model, comprising random walk and directed motion. For finding associations between the predictions and the detections at subsequent time points, a multi-frame approach was used, which is based on a graph theoretical formulation and supports one-to-one associations, many-to-one and one-to-many associations. The optimal associations were computed by solving a linear program (Jaiswal et al., 2015). The state of vesicles was computed by the Kalman filter based on the resulting associations and the selected motion model.

### Data processing and analysis

For image processing and analysis, the open-source image analysis software Fiji was used. Time-lapse images were registered in xy using the Fiji plugin “moco” (Dubbs et al., 2016). Average intensity projections of time-lapses was used to show stationary cell organelles as well as the outlines of axons, while maximum intensity projection was used to identify axons displaying active cell organelle trafficking. For mitochondrial transport, only active axonal stretches (ROIs) for analysis were selected. A total of 15 ROIs was analyzed per focal plane. Kymographs were generated with the Multi Kymograph tool (line width = 3) in Fiji. Kymographs revealed the number of stationary and mobile mitochondria, from which a mobility ratio was calculated. Kymographs were used to calculate the mean velocity of a mitochondrial track by using a Fiji macro written by Alessandro Moro. DCV transport was automatically tracked (see next section) and analyzed using self-written routines in Matlab (MathWorks). For mobility, DCV tracks were defined as mobile, if the total movement distance was above or equal to 5μm and stable if the movement distance was below 5μm. Mobility was calculated as mobile-fraction / (stable fraction + mobile fraction). Mean velocity was calculated on the mobile fraction only.

The mitoGFP and SyPhy-mCherry 3D-stacks were registered by manual selection of corresponding points and calculation of the resulting transformation in Matlab. For analyzing the structural plasticity of mitochondria and presynaptic terminals axons clearly visible in all time points (pre, 4 h and 24 h post) were chosen. Signals 1.5 times higher than the background of the axon were counted as mitochondria or presynaptic terminals, respectively.

### Statistical analysis

Statistical analysis was performed in Prism (GraphPad Software). For parametric tests, normal distribution was tested using D’Agostino & Pearson normality test. For analysis either goodness of fit, Mann-Whitney test, two-way repeated measures ANOVA, or Kruskal-Wallis were performed. For avoiding α-error accumulation, results were adjusted using post-hoc Sidak’s multiple comparisons test (two way repeated-measures ANOVAs) or Dunn’s multiple comparison test (Kruskal-Wallis). F values as well as degrees of freedom (DF; F (DFn, DFd)) are provided for ANOVAs. Statistical significance was assumed for P < 0.05. All data are displayed as mean ± standard deviation (STD) if not indicated otherwise (e.g. as mean ± standard error of the mean (SEM)).

### Behavioral testing in mice - Go/NoGo task

The Go/No-go task assesses the ability of mice to appropriately perform or withhold a lever pressing depending on the cues presented. Mice were trained for the Go/NoGo task in six operant chambers (TSE Systems). Each chamber was equipped with two ultrasensitive levers (required force, 1 g) on opponent sides: one functioning as the active and one as the inactive lever. Next to each lever, a front panel containing the visual stimulus was installed above a drinking microreservoir. When the programmed ratio requirements were met on the active lever, 10 μl of the sucrose (5%) solution was delivered into the microreservoir, and the visual stimulus was presented via a light located on the front panel. Responses on the inactive lever were recorded but had no programmed consequences. These responses were recorded as a measure of nonspecific behavioral activation. A microcomputer controlled the delivery of fluids, presentation of visual stimuli, and recording of the behavioral data.

The Go/No-go task was adapted from Gubner and colleagues (Gubner et al., 2010) and consisted of two training phases and one experimental phase. Briefly during each trial there was a variable duration pre-cue period (9-24 seconds) during which the house light was illuminated followed by a 5-seconds cue period, where one of two distinct cues (light of different colors in opposite walls) was used to differentiate Go trials from No-go trials. During a Go trial, a lever pressing terminated the Go cue and was reinforced with sucrose solution. Mice were permitted 3 s to consume the reinforcer, and then the 10-s inter-trial interval (ITI) began, during which the house light was off. During a No-go trial, a no response was reinforced followed by the 3-s consumption period and a 10-s ITI.

We measured the following variables: the active and inactive lever pressings, the pre-cue period duration, total active pressings during the pre-cue phase, number of right responses to go (correct go), number of missing the chances when they should go (missed go), number of right responses of no go (correct NoGo), number of going when they should not go (false alarm), efficiency (total number of reinforcers earned/total number of lever presses), pre-cue response rate, number of correct go in all sessions (go rate) and learning performance index (correct go/active lever pressings). False alarms and pre-cue response rates were the two main dependent measures of behavioral inhibition for the Go/No-go task.

During the experimental phase, i.e. after achieving a stable performance, mice were injected intraperitoneally (i.p.) with appropriate volumes of a 20% (v/v) ethanol solution to attain a dose of 3.5 g/kg body weight and confined to the operant boxes 4-6 h later. To test long-lasting effect, the test was repeated 24 and 48h later.

#### Statistics

Statistical analyses were performed by Student’s t test using Statistica 10 (StatSoft). All values are given as mean ± STD, and statistical significance was set at *p* < 0.05.

### *Drosophila* experiments

#### Fly strains

The following fly lines have been used: *TH*-Gal4 (Friggi-Grelin et al., 2003); *UAS-milton-RNAi^GD8116^*, *UAS-milton-RNAi^TRiPJF03022^* and *UAS-dmiro-RNAi^TRiPJF02775^*. RNAi–mediated knockdown of *milton-RN4i^TRiPJF03022^* reduced milton RNA levels and changed the axonal distribution of mitochondria similar to knock downs using the *UAS-milton-RNAi^GD8116^* and *UAS-dmiro-RNAi^TRiPJF02775^* showing that the transgenes are functional (Iijima-Ando et al., 2012).

For every experiment, 35 virgins were crossed to 15 male flies and raised on an ethanol-free standard cornmeal/molasses/yeast/agar medium on 12-h/12-h light/dark cycle at 25°C with 60% humidity. The transgenes were crossed to *w^1118^* to control for putative effect of the P-element insertion site of the transgene. The *TH*-Gal4 and *UAS-milton-RNAi^GD8116^* transgenes were at least backcrossed for 5 generation to the *w^1118^* stock of the Scholz lab to isogenize genetic background. To perform learning and memory experiments 50 zero to one day old male flies were collected under CO_2_ sedation and for recovery of sedation placed for 2 days on 25°C.

#### Conditioned preference for an odor associated with ethanol

The olfactory conditioned preference was performed according to (Nunez et al., 2018). Briefly a population of 50 flies were exposed to odor A (IA) in humidified air for 10 min followed by a second exposure to odor B (EA/AA) reinforced with ethanol for 10 min. After 50 min rest flies were trained again. The animals were trained in three cycles and the memory was tested 30 min later in a T-maze with a binary choice between odor A and odor B. The training was also done reciprocally with a different group of flies. The odorant/odor was diluted in mineral oil. The dilution for IA (VWR Life Science, 0944-1L) was 1:36 and the dilutions for EA1:36 and AA 1: 400 were mixed in a 2:1 ratio of EA (Sigma-Aldrich, 58958-5ML) and AA (Sigma-Aldrich, 71251-5ML-F). The ethanol vapor was generated by bubbling 95% EtOH. The vaporized EtOH was combined with air that was streamed over the odor cups. The flow rate was 294 U for vaporized ethanol and 117 U for humidified air (where 100 U is equal to 1.7 l min-1 at 20°C) to get an approximately 70% Ethanol ratio in the air.

The preference index was calculated as follows: (number of flies in paired odor vial – number of flies in unpaired odor vial) / total number of flies. The average of the preference indexes of two reciprocally trained fly groups is used as conditioned odor preference or aversion (CPI). A positive CPI indicates a positive association with ethanol for the animals.

The sensory acuity was performed by exposing flies in a T-maze under similar condition as the test phase of the learning and memory experiment. However here flies were not previous exposed to an odor/odorant.

#### Statistics

The One-sample sign test was used to determine difference from random choice and significant differences with *P* < 0.05 were indicated with the letter “**a**”. Differences between more than two test groups were determined using ANOVA *post-hoc* Tukey-Kramer Honestly Significant Difference (HSD). *P\** < 0.05, *P\*\** < 0.01 and *P\*\*\** < 0.001. The errors are given as s.e.m..

## Notes

https://heidata.uni-heidelberg.de/privateurl.xhtml?token=f88c8325-6f37-473d-856a-0af8109a2bba

### Competing Interest Statement

The authors have declared no competing interest.

## References

Alia-Klein, N., Goldstein, R.Z., Kriplani, A., Logan, J., Tomasi, D., Williams, B., Telang, F., Shumay, E., Biegon, A., Craig, I.W., et al. (2008). Brain monoamine oxidase A activity predicts trait aggression. J Neurosci 28, 5099–5104.

Andrieux, A., Salin, P.A., Vernet, M., Kujala, P., Baratier, J., Gory-Faure, S., Bosc, C., Pointu, H., Proietto, D., Schweitzer, A., et al. (2002). The suppression of brain cold-stable microtubules in mice induces synaptic defects associated with neuroleptic-sensitive behavioral disorders. Genes Dev 16, 2350–2364.

Ash, R.T., Fahey, P.G., Park, J., Zoghbi, H.Y., and Smirnakis, S.M. (2018). Increased Axonal Bouton Stability during Learning in the Mouse Model of MECP2 Duplication Syndrome. eNeuro 5.

Berthet, A., Margolis, E.B., Zhang, J., Hsieh, I., Zhang, J., Hnasko, T.S., Ahmad, J., Edwards, R.H., Sesaki, H., Huang, E.J., et al. (2014). Loss of mitochondrial fission depletes axonal mitochondria in midbrain dopamine neurons. J Neurosci 34, 14304–14317.

Boffi, J.C., Knabbe, J., Kaiser, M., and Kuner, T. (2018). KCC2-dependent Steady-state Intracellular Chloride Concentration and pH in Cortical Layer 2/3 Neurons of Anesthetized and Awake Mice. Front Cell Neurosci 12, 7.

Bukalo, O., Schachner, M., and Dityatev, A. (2007). Hippocampal metaplasticity induced by deficiency in the extracellular matrix glycoprotein tenascin-R. J Neurosci 27, 6019–6028.

Camarini, R., and Hodge, C.W. (2004). Ethanol preexposure increases ethanol self-administration in C57BL/6J and DBA/2J mice. Pharmacol Biochem Behav 79, 623–632.

Carpenter-Hyland, E.P., and Chandler, L.J. (2006). Homeostatic plasticity during alcohol exposure promotes enlargement of dendritic spines. Eur J Neurosci 24, 3496–3506.

Cheng, D., Hoogenraad, C.C., Rush, J., Ramm, E., Schlager, M.A., Duong, D.M., Xu, P., Wijayawardana, S.R., Hanfelt, J., Nakagawa, T., et al. (2006). Relative and absolute quantification of postsynaptic density proteome isolated from rat forebrain and cerebellum. Mol Cell Proteomics 5, 1158–1170.

Ciccocioppo, R., Martin-Fardon, R., and Weiss, F. (2004). Stimuli associated with a single cocaine experience elicit long-lasting cocaine-seeking. Nature neuroscience 7, 495–496.

Conigrave, K.M., Davies, P., Haber, P., and Whitfield, J.B. (2003). Traditional markers of excessive alcohol use. Addiction 98 Suppl 2, 31–43.

Cox, J.M., M (2008). High peptide identification rates, individualized ppb-range mass accuracies and proteome-wide quantitation via novel computational strategies. in 2nd revision. Nature Biotechnology.

Cui, H., Darmanin, S., Natsuisaka, M., Kondo, T., Asaka, M., Shindoh, M., Higashino, F., Hamuro, J., Okada, F., Kobayashi, M., et al. (2007). Enhanced expression of asparagine synthetase under glucose-deprived conditions protects pancreatic cancer cells from apoptosis induced by glucose deprivation and cisplatin. Cancer research 67, 3345–3355.

de Wit, J., Toonen, R.F., Verhaagen, J., and Verhage, M. (2006). Vesicular trafficking of semaphorin 3A is activity-dependent and differs between axons and dendrites. Traffic 7, 1060–1077.

Denk, W., Delaney, K.R., Gelperin, A., Kleinfeld, D., Strowbridge, B.W., Tank, D.W., and Yuste, R. (1994). Anatomical and functional imaging of neurons using 2-photon laser scanning microscopy. J Neurosci Methods 54, 151–162.

DeWit, D.J., Adlaf, E.M., Offord, D.R., and Ogborne, A.C. (2000). Age at first alcohol use: a risk factor for the development of alcohol disorders. Am J Psychiatry 157, 745–750.

Dubbs, A., Guevara, J., and Yuste, R. (2016). moco: Fast Motion Correction for Calcium Imaging. Front Neuroinform 10, 6.

Eisenhardt, M., Hansson, A.C., Spanagel, R., and Bilbao, A. (2015). Chronic intermittent ethanol exposure in mice leads to an up-regulation of CRH/CRHR1 signaling. Alcohol Clin Exp Res 39, 752–762.

Engert, F., and Bonhoeffer, T. (1999). Dendritic spine changes associated with hippocampal long-term synaptic plasticity. Nature 399, 66–70.

Ernst, L., Darschnik, S., Roos, J., Gonzalez-Gomez, M., Beemelmans, C., Beemelmans, C., Engelhardt, M., Meyer, G., and Wahle, P. (2018). Fast prenatal development of the NPY neuron system in the neocortex of the European wild boar, Sus scrofa. Brain Struct Funct 223, 3855–3873.

Evans, M.D., Dumitrescu, A.S., Kruijssen, D.L.H., Taylor, S.E., and Grubb, M.S. (2015). Rapid Modulation of Axon Initial Segment Length Influences Repetitive Spike Firing. Cell Rep 13, 1233–1245.

Faingold, C.L., N’Gouemo, P., and Riaz, A. (1998). Ethanol and neurotransmitter interactions--from molecular to integrative effects. Prog Neurobiol 55, 509–535.

Feng, G., Mellor, R.H., Bernstein, M., Keller-Peck, C., Nguyen, Q.T., Wallace, M., Nerbonne, J.M., Lichtman, J.W., and Sanes, J.R. (2000). Imaging neuronal subsets in transgenic mice expressing multiple spectral variants of GFP. Neuron 28, 41–51.

Ferrandina, G., Zannoni, G.F., Martinelli, E., Paglia, A., Gallotta, V., Mozzetti, S., Scambia, G., and Ferlini, C. (2006). Class III beta-tubulin overexpression is a marker of poor clinical outcome in advanced ovarian cancer patients. Clin Cancer Res 12, 2774–2779.

Fonseca, R., Vabulas, R.M., Hartl, F.U., Bonhoeffer, T., and Nagerl, U.V. (2006). A balance of protein synthesis and proteasome-dependent degradation determines the maintenance of LTP. Neuron 52, 239–245.

Forstera, B., Castro, P.A., Moraga-Cid, G., and Aguayo, L.G. (2016). Potentiation of Gamma Aminobutyric Acid Receptors (GABAAR) by Ethanol: How Are Inhibitory Receptors Affected? Front Cell Neurosci 10, 114.

Frank, A.C., Huang, S., Zhou, M., Gdalyahu, A., Kastellakis, G., Silva, T.K., Lu, E., Wen, X., Poirazi, P., Trachtenberg, J.T., et al. (2018). Hotspots of dendritic spine turnover facilitate clustered spine addition and learning and memory. Nat Commun 9, 422.

Friggi-Grelin, F., Coulom, H., Meller, M., Gomez, D., Hirsh, J., and Birman, S. (2003). Targeted gene expression in Drosophila dopaminergic cells using regulatory sequences from tyrosine hydroxylase. Journal of neurobiology 54, 618–627.

Fullgrabe, M.W., Vengeliene, V., and Spanagel, R. (2007). Influence of age at drinking onset on the alcohol deprivation effect and stress-induced drinking in female rats. Pharmacol Biochem Behav 86, 320–326.

Ginsburg, B.C., Javors, M.A., Friesenhahn, G., Frontz, M., Martinez, G., Hite, T., and Lamb, R.J. (2008). Mouse breathalyzer. Alcohol Clin Exp Res 32, 1181–1185.

Glater, E.E., Megeath, L.J., Stowers, R.S., and Schwarz, T.L. (2006). Axonal transport of mitochondria requires milton to recruit kinesin heavy chain and is light chain independent. The Journal of cell biology 173, 545–557.

Grant, S.G.N. (2018). Synapse molecular complexity and the plasticity behaviour problem. Brain Neurosci Adv 2, 2398212818810685.

Grubb, M.S., and Burrone, J. (2010). Activity-dependent relocation of the axon initial segment fine-tunes neuronal excitability. Nature 465, 1070–1074.

Gubner, N.R., Wilhelm, C.J., Phillips, T.J., and Mitchell, S.H. (2010). Strain differences in behavioral inhibition in a Go/No-go task demonstrated using 15 inbred mouse strains. Alcohol Clin Exp Res 34, 1353–1362.

Guo, A.Y., Webb, B.T., Miles, M.F., Zimmerman, M.P., Kendler, K.S., and Zhao, Z. (2009). ERGR: An ethanol-related gene resource. Nucleic Acids Res 37, D840–845.

Guo, X., Macleod, G.T., Wellington, A., Hu, F., Panchumarthi, S., Schoenfield, M., Marin, L., Charlton, M.P., Atwood, H.L., and Zinsmaier, K.E. (2005). The GTPase dMiro is required for axonal transport of mitochondria to Drosophila synapses. Neuron 47, 379–393.

Halbout, B., Bernardi, R.E., Hansson, A.C., and Spanagel, R. (2014). Incubation of cocaine seeking following brief cocaine experience in mice is enhanced by mGluR1 blockade. J Neurosci 34, 1781–1790.

Haseba, T., Kameyama, K., Mashimo, K., and Ohno, Y. (2012). Dose-Dependent Change in Elimination Kinetics of Ethanol due to Shift of Dominant Metabolizing Enzyme from ADH 1 (Class I) to ADH 3 (Class III) in Mouse. Int J Hepatol 2012, 408190.

Hayashi, Y., Shi, S.H., Esteban, J.A., Piccini, A., Poncer, J.C., and Malinow, R. (2000). Driving AMPA receptors into synapses by LTP and CaMKII: requirement for GluR1 and PDZ domain interaction. Science 287, 2262–2267.

Heinz, A., Kiefer, F., Smolka, M.N., Endrass, T., Beste, C., Beck, A., Liu, S., Genauck, A., Romund, L., Banaschewski, T., et al. (2020). Addiction Research Consortium: Losing and regaining control over drug intake (ReCoDe)-From trajectories to mechanisms and interventions. Addict Biol 25, e12866.

Henry, K.L., McDonald, J.N., Oetting, E.R., Walker, P.S., Walker, R.D., and Beauvais, F. (2011). Age of onset of first alcohol intoxication and subsequent alcohol use among urban American Indian adolescents. Psychol Addict Behav 25, 48–56.

Hingson, R., Heeren, T., Zakocs, R., Winter, M., and Wechsler, H. (2003). Age of first intoxication, heavy drinking, driving after drinking and risk of unintentional injury among U.S. college students. J Stud Alcohol 64, 23–31.

Hirth, N., Meinhardt, M.W., Noori, H.R., Salgado, H., Torres-Ramirez, O., Uhrig, S., Broccoli, L., Vengeliene, V., Rossmanith, M., Perreau-Lenz, S., et al. (2016). Convergent evidence from alcohol-dependent humans and rats for a hyperdopaminergic state in protracted abstinence. Proc Natl Acad Sci U S A 113, 3024–3029.

Hofflin, F., Jack, A., Riedel, C., Mack-Bucher, J., Roos, J., Corcelli, C., Schultz, C., Wahle, P., and Engelhardt, M. (2017). Heterogeneity of the Axon Initial Segment in Interneurons and Pyramidal Cells of Rodent Visual Cortex. Front Cell Neurosci 11, 332.

Holtmaat, A., Bonhoeffer, T., Chow, D.K., Chuckowree, J., De Paola, V., Hofer, S.B., Hubener, M., Keck, T., Knott, G., Lee, W.C., et al. (2009). Long-term, high-resolution imaging in the mouse neocortex through a chronic cranial window. Nat Protoc 4, 1128–1144.

Holtmaat, A.J., Trachtenberg, J.T., Wilbrecht, L., Shepherd, G.M., Zhang, X., Knott, G.W., and Svoboda, K. (2005). Transient and persistent dendritic spines in the neocortex in vivo. Neuron 45, 279–291.

Hou, Q.L., Gao, X., Lu, Q., Zhang, X.H., Tu, Y.Y., Jin, M.L., Zhao, G.P., Yu, L., Jing, N.H., and Li, B.M. (2006). SNAP-25 in hippocampal CA3 region is required for long-term memory formation. Biochem Biophys Res Commun 347, 955–962.

Hulo, S., Alberi, S., Laux, T., Muller, D., and Caroni, P. (2002). A point mutant of GAP-43 induces enhanced short-term and long-term hippocampal plasticity. Eur J Neurosci 15, 1976–1982.

Hyman, S.E., Malenka, R.C., and Nestler, E.J. (2006). Neural mechanisms of addiction: the role of reward-related learning and memory. Annu Rev Neurosci 29, 565–598.

Iijima-Ando, K., Sekiya, M., Maruko-Otake, A., Ohtake, Y., Suzuki, E., Lu, B., and Iijima, K.M. (2012). Loss of axonal mitochondria promotes tau-mediated neurodegeneration and Alzheimer’s disease-related tau phosphorylation via PAR-1. PLoS genetics 8, e1002918.

Inoue, E., Mochida, S., Takagi, H., Higa, S., Deguchi-Tawarada, M., Takao-Rikitsu, E., Inoue, M., Yao, I., Takeuchi, K., Kitajima, I., et al. (2006). SAD: a presynaptic kinase associated with synaptic vesicles and the active zone cytomatrix that regulates neurotransmitter release. Neuron 50, 261–275.

Isshiki, M., Tanaka, S., Kuriu, T., Tabuchi, K., Takumi, T., and Okabe, S. (2014). Enhanced synapse remodelling as a common phenotype in mouse models of autism. Nat Commun 5, 4742.

Jaiswal, A., Godinez, W.J., Eils, R., Lehmann, M.J., and Rohr, K. (2015). Tracking Virus Particles in Fluorescence Microscopy Images Using Multi-Scale Detection and Multi-Frame Association. IEEE Trans Image Process 24, 4122–4136.

Jamann, N., Jordan, M., and Engelhardt, M. (2018). Activity-dependent axonal plasticity in sensory systems. Neuroscience 368, 268–282.

Jin, W., Scotto, K.W., Hait, W.N., and Yang, J.M. (2007). Involvement of CtBP1 in the transcriptional activation of the MDR1 gene in human multidrug resistant cancer cells. Biochem Pharmacol 74, 851–859.

Kaech, S., and Banker, G. (2006). Culturing hippocampal neurons. Nat Protoc 1, 2406–2415.

Kalman, R.E. (1960). A new approach to linear filtering and prediction problems. J Basic Eng D 82, 35–45.

Kaun, K.R., Azanchi, R., Maung, Z., Hirsh, J., and Heberlein, U. (2011). A Drosophila model for alcohol reward. Nature neuroscience 14, 612–619.

Kim, H.J., Choi, K.M., Ku, B.M., Mun, J., Joo, Y., Han, J.Y., Kim, Y.H., Roh, G.S., Kang, S.S., Cho, G.J., et al. (2006). Acute ethanol administration decreases GAP-43 and phosphorylated-GAP-43 in the rat hippocampus. Brain Res 1112, 16–25.

Kiryu-Seo, S., Ohno, N., Kidd, G.J., Komuro, H., and Trapp, B.D. (2010). Demyelination increases axonal stationary mitochondrial size and the speed of axonal mitochondrial transport. J Neurosci 30, 6658–6666.

Knabbe, J., Nassal, J.P., Verhage, M., and Kuner, T. (2018). Secretory vesicle trafficking in awake and anaesthetized mice: differential speeds in axons versus synapses. J Physiol 596, 3759–3773.

Krantic, S., Mechawar, N., Reix, S., and Quirion, R. (2007). Apoptosis-inducing factor: a matter of neuron life and death. Prog Neurobiol 81, 179–196.

Kruger, M., Moser, M., Ussar, S., Thievessen, I., Luber, C.A., Forner, F., Schmidt, S., Zanivan, S., Fassler, R., and Mann, M. (2008). SILAC mouse for quantitative proteomics uncovers kindlin-3 as an essential factor for red blood cell function. Cell 134, 353–364.

Kwinter, D.M., Lo, K., Mafi, P., and Silverman, M.A. (2009). Dynactin regulates bidirectional transport of dense-core vesicles in the axon and dendrites of cultured hippocampal neurons. Neuroscience 162, 1001–1010.

Lebedeva, J., Zakharov, A., Ogievetsky, E., Minlebaeva, A., Kurbanov, R., Gerasimova, E., Sitdikova, G., and Khazipov, R. (2017). Inhibition of Cortical Activity and Apoptosis Caused by Ethanol in Neonatal Rats In Vivo. Cereb Cortex 27, 1068–1082.

Lein, P.J., Barnhart, C.D., and Pessah, I.N. (2011). Acute hippocampal slice preparation and hippocampal slice cultures. Methods Mol Biol 758, 115–134.

Leterrier, C. (2018). The Axon Initial Segment: An Updated Viewpoint. J Neurosci 38, 2135–2145.

Lewis, T.L., Jr., Turi, G.F., Kwon, S.K., Losonczy, A., and Polleux, F. (2016). Progressive Decrease of Mitochondrial Motility during Maturation of Cortical Axons In Vitro and In Vivo. Curr Biol 26, 2602–2608.

Miyakawa, T., Yagi, T., Kitazawa, H., Yasuda, M., Kawai, N., Tsuboi, K., and Niki, H. (1997). Fyn-kinase as a determinant of ethanol sensitivity: relation to NMDA-receptor function. Science 278, 698–701.

Morean, M.E., Kong, G., Camenga, D.R., Cavallo, D.A., Connell, C., and Krishnan-Sarin, S. (2014). First drink to first drunk: age of onset and delay to intoxication are associated with adolescent alcohol use and binge drinking. Alcohol Clin Exp Res 38, 2615–2621.

Nagerl, U.V., Eberhorn, N., Cambridge, S.B., and Bonhoeffer, T. (2004). Bidirectional activity-dependent morphological plasticity in hippocampal neurons. Neuron 44, 759–767.

Nonn, L., Williams, R.R., Erickson, R.P., and Powis, G. (2003). The absence of mitochondrial thioredoxin 2 causes massive apoptosis, exencephaly, and early embryonic lethality in homozygous mice. Mol Cell Biol 23, 916–922.

Nunez, K.M., Azanchi, R., and Kaun, K.R. (2018). Cue-Induced Ethanol Seeking in Drosophila melanogaster Is Dose-Dependent. Front Physiol 9, 438.

Okabe, S. (2017). Fluorescence imaging of synapse dynamics in normal circuit maturation and in developmental disorders. Proc Jpn Acad Ser B Phys Biol Sci 93, 483–497.

Ong, S.E., Foster, L.J., and Mann, M. (2003). Mass spectrometric-based approaches in quantitative proteomics. Methods 29, 124–130.

Ou, X.M., Jafar-Nejad, H., Storring, J.M., Meng, J.H., Lemonde, S., and Albert, P.R. (2000). Novel dual repressor elements for neuronal cell-specific transcription of the rat 5-HT1A receptor gene. J Biol Chem 275, 8161–8168.

Pandya, N.J., Koopmans, F., Slotman, J.A., Paliukhovich, I., Houtsmuller, A.B., Smit, A.B., and Li, K.W. (2017). Correlation profiling of brain sub-cellular proteomes reveals co-assembly of synaptic proteins and subcellular distribution. Sci Rep 7, 12107.

Papa, M., and Segal, M. (1996). Morphological plasticity in dendritic spines of cultured hippocampal neurons. Neuroscience 71, 1005–1011.

Patel, V.B., Sandhu, G., Corbett, J.M., Dunn, M.J., Rodrigues, L.M., Griffiths, J.R., Wassif, W., Sherwood, R.A., Richardson, P.J., and Preedy, V.R. (2000). A comparative investigation into the effect of chronic alcohol feeding on the myocardium of normotensive and hypertensive rats: an electrophoretic and biochemical study. Electrophoresis 21, 2454–2462.

Peris, L., Bisbal, M., Martinez-Hernandez, J., Saoudi, Y., Jonckheere, J., Rolland, M., Sebastien, M., Brocard, J., Denarier, E., Bosc, C., et al. (2018). A key function for microtubule-associated-protein 6 in activity-dependent stabilisation of actin filaments in dendritic spines. Nat Commun 9, 3775.

Phillips, G.R., Huang, J.K., Wang, Y., Tanaka, H., Shapiro, L., Zhang, W., Shan, W.S., Arndt, K., Frank, M., Gordon, R.E., et al. (2001). The presynaptic particle web: ultrastructure, composition, dissolution, and reconstitution. Neuron 32, 63–77.

Pilling, A.D., Horiuchi, D., Lively, C.M., and Saxton, W.M. (2006). Kinesin-1 and Dynein are the primary motors for fast transport of mitochondria in Drosophila motor axons. Molecular biology of the cell 17, 2057–2068.

Poo, M.M., Pignatelli, M., Ryan, T.J., Tonegawa, S., Bonhoeffer, T., Martin, K.C., Rudenko, A., Tsai, L.H., Tsien, R.W., Fishell, G., et al. (2016). What is memory? The present state of the engram. BMC Biol 14, 40.

Popova, N.K., Vishnivetskaya, G.B., Ivanova, E.A., Skrinskaya, J.A., and Seif, I. (2000). Altered behavior and alcohol tolerance in transgenic mice lacking MAO A: a comparison with effects of MAO A inhibitor clorgyline. Pharmacol Biochem Behav 67, 719–727.

Roberts, T.F., Tschida, K.A., Klein, M.E., and Mooney, R. (2010). Rapid spine stabilization and synaptic enhancement at the onset of behavioural learning. Nature 463, 948–952.

Rogaeva, A., Ou, X.M., Jafar-Nejad, H., Lemonde, S., and Albert, P.R. (2007). Differential repression by freud-1/CC2D1A at a polymorphic site in the dopamine-D2 receptor gene. J Biol Chem 282, 20897–20905.

Schluter, A., Del Turco, D., Deller, T., Gutzmann, A., Schultz, C., and Engelhardt, M. (2017). Structural Plasticity of Synaptopodin in the Axon Initial Segment during Visual Cortex Development. Cereb Cortex 27, 4662–4675.

Schwenger, D.B., and Kuner, T. (2010). Acute genetic perturbation of exocyst function in the rat calyx of Held impedes structural maturation, but spares synaptic transmission. Eur J Neurosci 32, 974–984.

Sherif, F.M., Tawati, A.M., Ahmed, S.S., and Sharif, S.I. (1997). Basic aspects of GABA-transmission in alcoholism, with particular reference to GABA-transaminase. Eur Neuropsychopharmacol 7, 1–7.

Silva, J.C., Gorenstein, M.V., Li, G.Z., Vissers, J.P., and Geromanos, S.J. (2006). Absolute quantification of proteins by LCMSE: a virtue of parallel MS acquisition. Mol Cell Proteomics 5, 144–156.

Smit-Rigter, L., Rajendran, R., Silva, C.A., Spierenburg, L., Groeneweg, F., Ruimschotel, E.M., van Versendaal, D., van der Togt, C., Eysel, U.T., Heimel, J.A., et al. (2016). Mitochondrial Dynamics in Visual Cortex Are Limited In Vivo and Not Affected by Axonal Structural Plasticity. Curr Biol 26, 2609–2616.

Smith, H.L., Bourne, J.N., Cao, G., Chirillo, M.A., Ostroff, L.E., Watson, D.J., and Harris, K.M. (2016). Mitochondrial support of persistent presynaptic vesicle mobilization with age-dependent synaptic growth after LTP. Elife 5.

Smith, K.R., Kopeikina, K.J., Fawcett-Patel, J.M., Leaderbrand, K., Gao, R., Schurmann, B., Myczek, K., Radulovic, J., Swanson, G.T., and Penzes, P. (2014). Psychiatric risk factor ANK3/ankyrin-G nanodomains regulate the structure and function of glutamatergic synapses. Neuron 84, 399–415.

Spanagel, R., and Kiefer, F. (2008). Drugs for relapse prevention of alcoholism: ten years of progress. Trends Pharmacol Sci 29, 109–115.

Stowers, R.S., Megeath, L.J., Gorska-Andrzejak, J., Meinertzhagen, I.A., and Schwarz, T.L. (2002). Axonal transport of mitochondria to synapses depends on milton, a novel Drosophila protein. Neuron 36, 1063–1077.

Strack, S., Choi, S., Lovinger, D.M., and Colbran, R.J. (1997). Translocation of autophosphorylated calcium/calmodulin-dependent protein kinase II to the postsynaptic density. J Biol Chem 272, 13467–13470.

Sugino, K., Clark, E., Schulmann, A., Shima, Y., Wang, L., Hunt, D.L., Hooks, B.M., Trankner, D., Chandrashekar, J., Picard, S., et al. (2019). Mapping the transcriptional diversity of genetically and anatomically defined cell populations in the mouse brain. Elife 8.

Takeda-Uchimura, Y., Uchimura, K., Sugimura, T., Yanagawa, Y., Kawasaki, T., Komatsu, Y., and Kadomatsu, K. (2015). Requirement of keratan sulfate proteoglycan phosphacan with a specific sulfation pattern for critical period plasticity in the visual cortex. Exp Neurol 274, 145–155.

Takihara, Y., Inatani, M., Eto, K., Inoue, T., Kreymerman, A., Miyake, S., Ueno, S., Nagaya, M., Nakanishi, A., Iwao, K., et al. (2015). In vivo imaging of axonal transport of mitochondria in the diseased and aged mammalian CNS. Proc Natl Acad Sci U S A 112, 10515–10520.

Todorova, V., and Blokland, A. (2017). Mitochondria and Synaptic Plasticity in the Mature and Aging Nervous System. Curr Neuropharmacol 15, 166–173.

Tu, Y., Kroener, S., Abernathy, K., Lapish, C., Seamans, J., Chandler, L.J., and Woodward, J.J. (2007). Ethanol inhibits persistent activity in prefrontal cortical neurons. J Neurosci 27, 4765–4775.

Turrigiano, G. (2012). Homeostatic synaptic plasticity: local and global mechanisms for stabilizing neuronal function. Cold Spring Harb Perspect Biol 4, a005736.

Vaccaro, V., Devine, M.J., Higgs, N.F., and Kittler, J.T. (2017). Miro1-dependent mitochondrial positioning drives the rescaling of presynaptic Ca2+ signals during homeostatic plasticity. EMBO Rep 18, 231–240.

Van Skike, C.E., Goodlett, C., and Matthews, D.B. (2019). Acute alcohol and cognition: Remembering what it causes us to forget. Alcohol 79, 105–125.

Verstreken, P., Ly, C.V., Venken, K.J., Koh, T.W., Zhou, Y., and Bellen, H.J. (2005). Synaptic mitochondria are critical for mobilization of reserve pool vesicles at Drosophila neuromuscular junctions. Neuron 47, 365–378.

Wanat, M.J., Willuhn, I., Clark, J.J., and Phillips, P.E. (2009). Phasic dopamine release in appetitive behaviors and drug addiction. Curr Drug Abuse Rev 2, 195–213.

Wang, C., Zhou, G.L., Vedantam, S., Li, P., and Field, J. (2008). Mitochondrial shuttling of CAP1 promotes actin- and cofilin-dependent apoptosis. J Cell Sci 121, 2913–2920.

Wang, J., Carnicella, S., Phamluong, K., Jeanblanc, J., Ronesi, J.A., Chaudhri, N., Janak, P.H., Lovinger, D.M., and Ron, D. (2007). Ethanol induces long-term facilitation of NR2B-NMDA receptor activity in the dorsal striatum: implications for alcohol drinking behavior. J Neurosci 27, 3593–3602.

Weiner, J.L., and Valenzuela, C.F. (2006). Ethanol modulation of GABAergic transmission: the view from the slice. Pharmacol Ther 111, 533–554.

Whelan, R., Watts, R., Orr, C.A., Althoff, R.R., Artiges, E., Banaschewski, T., Barker, G.J., Bokde, A.L., Buchel, C., Carvalho, F.M., et al. (2014). Neuropsychosocial profiles of current and future adolescent alcohol misusers. Nature 512, 185–189.

White, A.M., Matthews, D.B., and Best, P.J. (2000). Ethanol, memory, and hippocampal function: a review of recent findings. Hippocampus 10, 88–93.

White, A.M., and Swartzwelder, H.S. (2004). Hippocampal function during adolescence: a unique target of ethanol effects. Ann N Y Acad Sci 1021, 206–220.

Wilhelm, B.G., Mandad, S., Truckenbrodt, S., Krohnert, K., Schafer, C., Rammner, B., Koo, S.J., Classen, G.A., Krauss, M., Haucke, V., et al. (2014). Composition of isolated synaptic boutons reveals the amounts of vesicle trafficking proteins. Science 344, 1023–1028.

Yin, J., and Yuan, Q. (2014). Structural homeostasis in the nervous system: a balancing act for wiring plasticity and stability. Front Cell Neurosci 8, 439.

Zahn, T.R., Angleson, J.K., MacMorris, M.A., Domke, E., Hutton, J.F., Schwartz, C., and Hutton, J.C. (2004). Dense core vesicle dynamics in Caenorhabditis elegans neurons and the role of kinesin UNC-104. Traffic 5, 544–559.

Zhang, H.T., Huang, Y., Jin, S.L., Frith, S.A., Suvarna, N., Conti, M., and O’Donnell, J.M. (2002). Antidepressant-like profile and reduced sensitivity to rolipram in mice deficient in the PDE4D phosphodiesterase enzyme. Neuropsychopharmacology 27, 587–595.

