## Supplementary figures and images for "Single-dose ethanol intoxication causes acute and lasting neuronal changes in the brain"

### Supplemental Figure 1

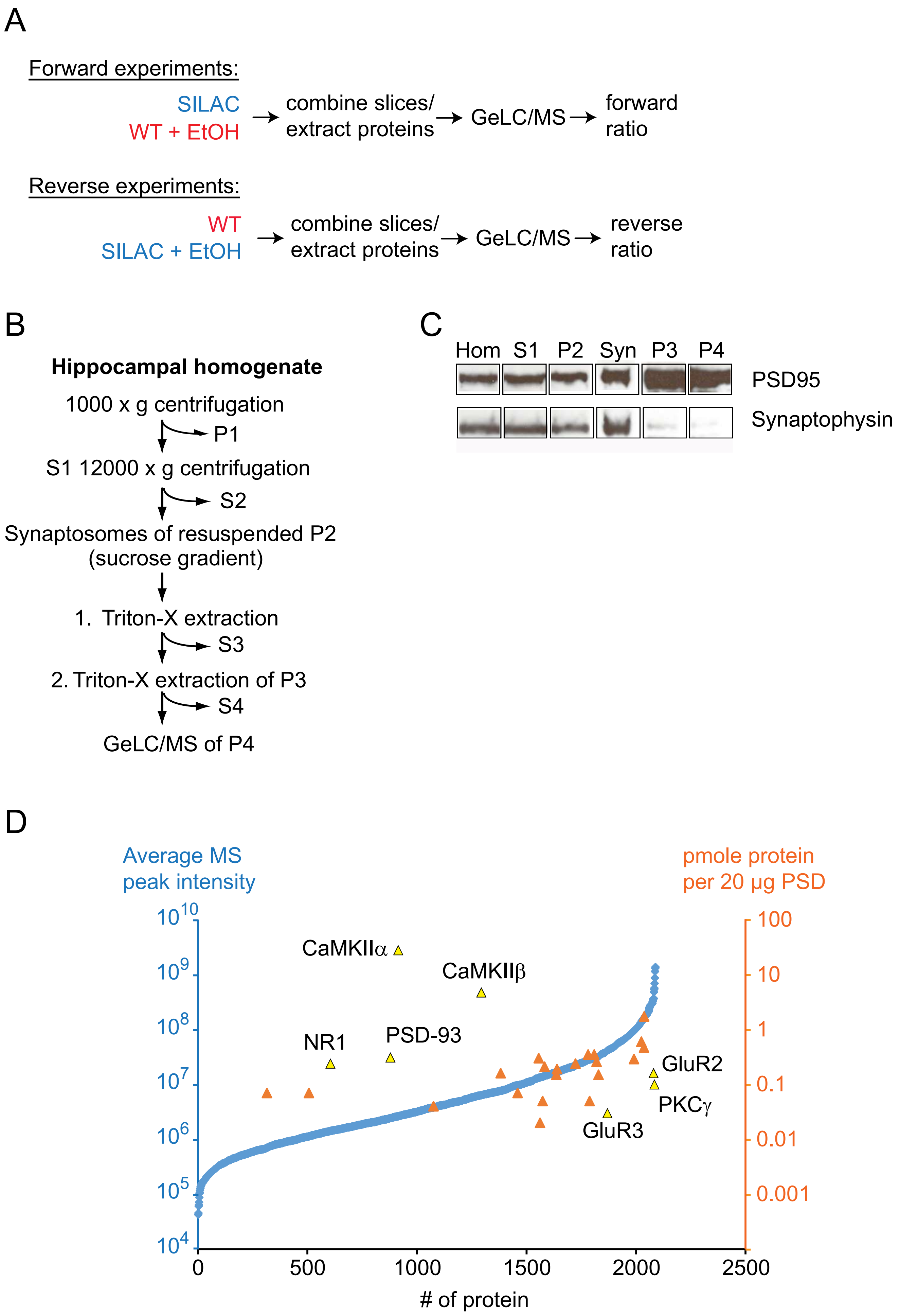


#### Supplementary Figure 1

### Supplemental Figure 2

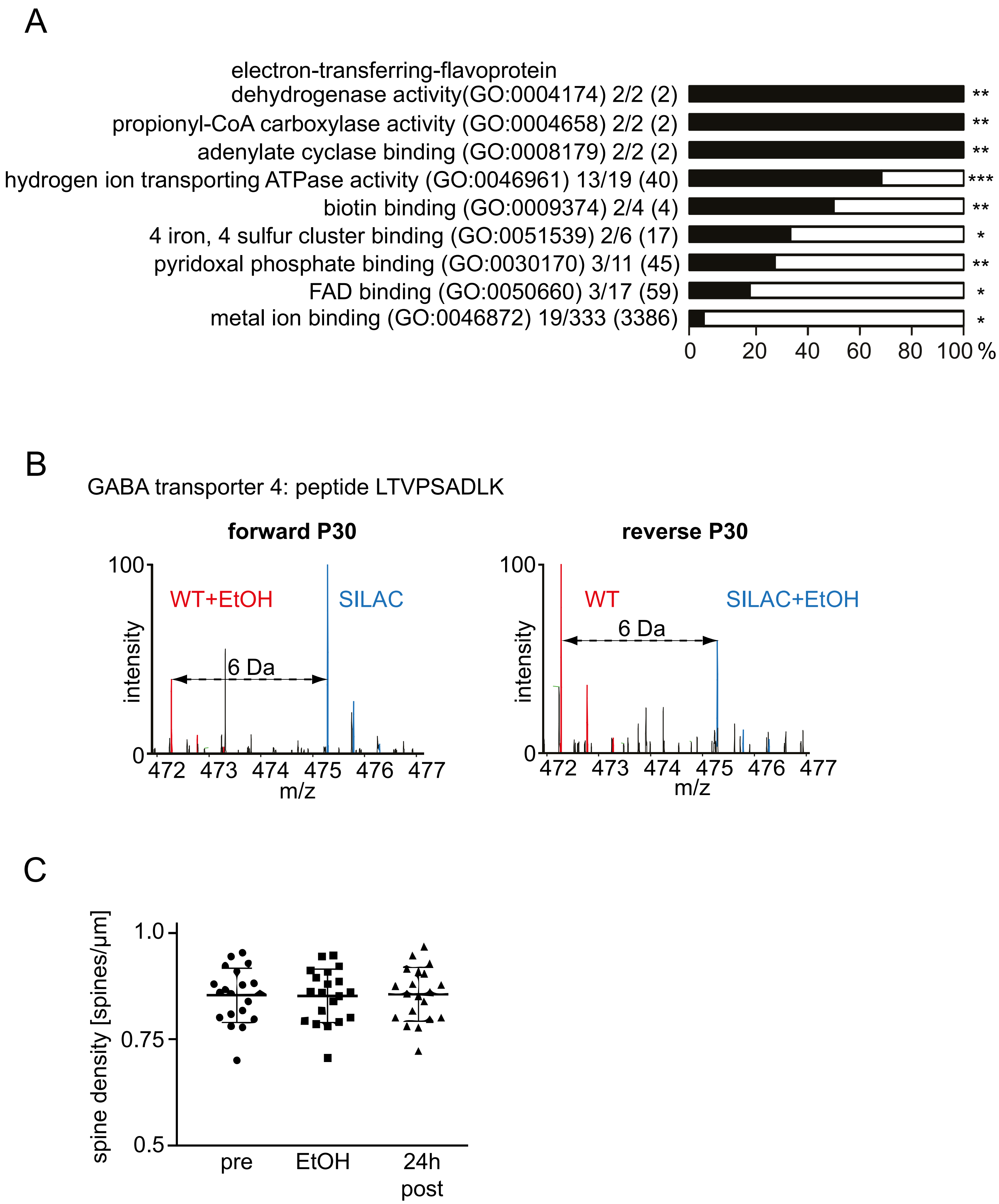


Supplementary Figure 2

### Supplemental Figure 3

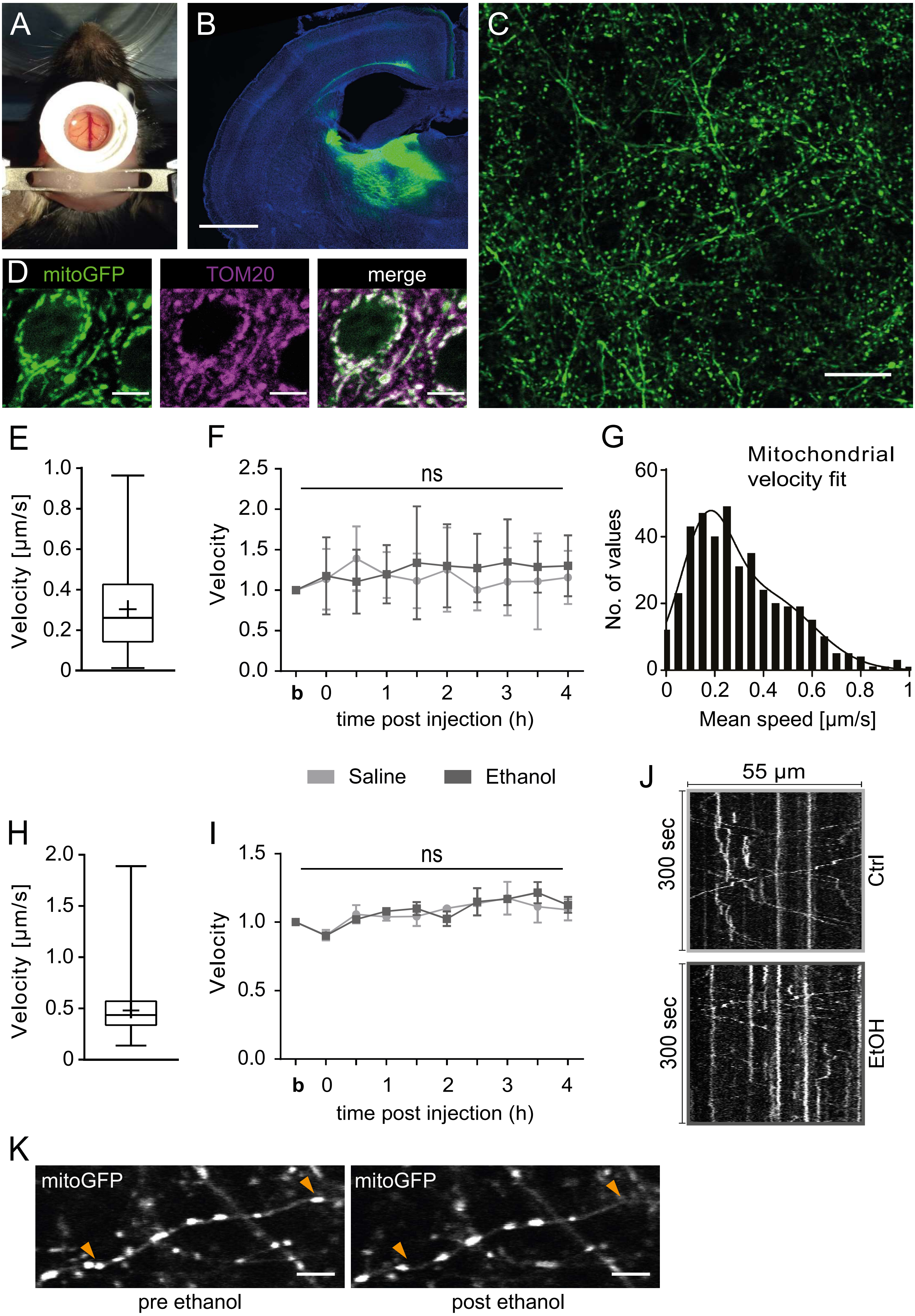


Supplementary Figure 3

### Supplemental Figure 4

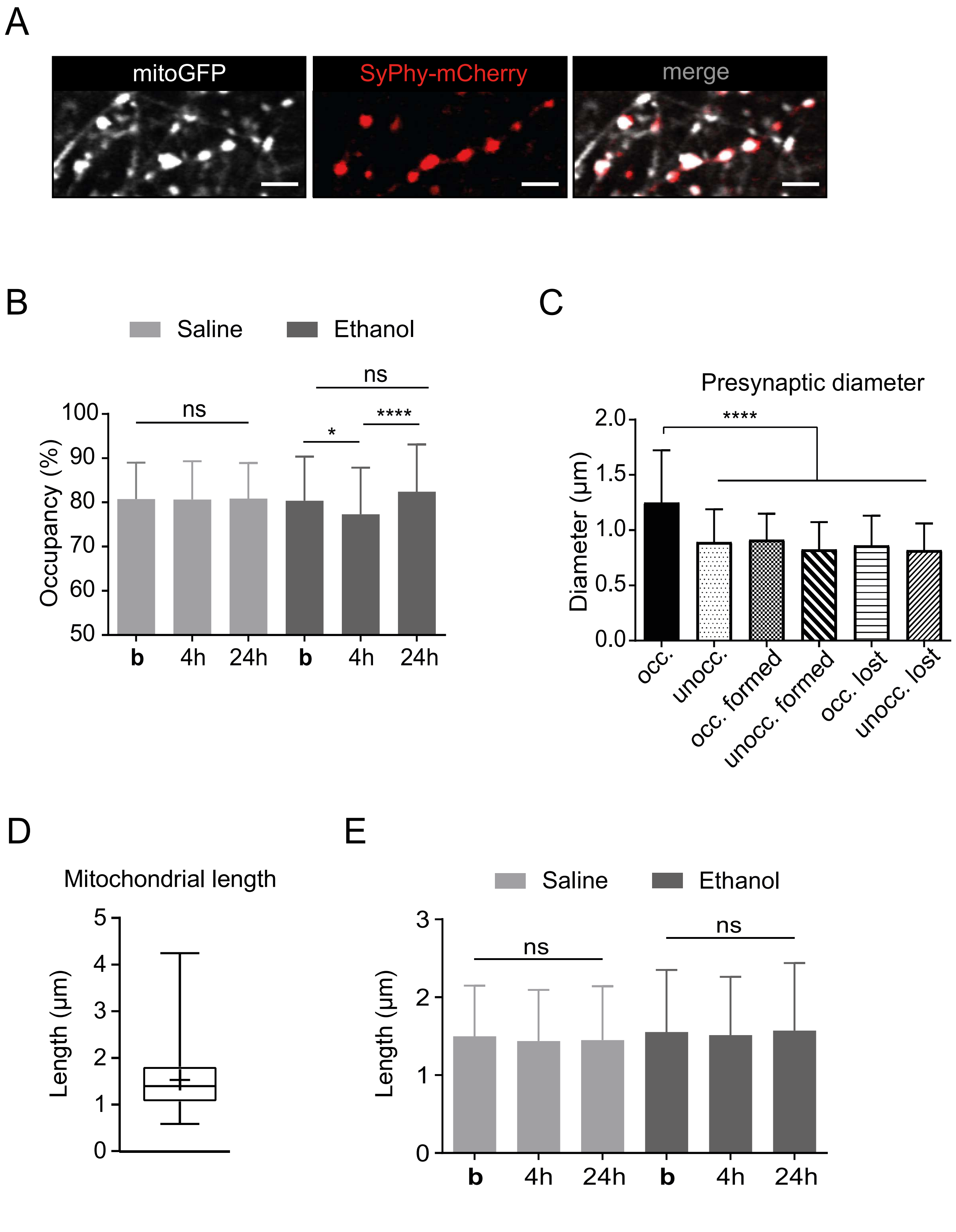


Supplementary Figure 4

### Supplemental Figure 5

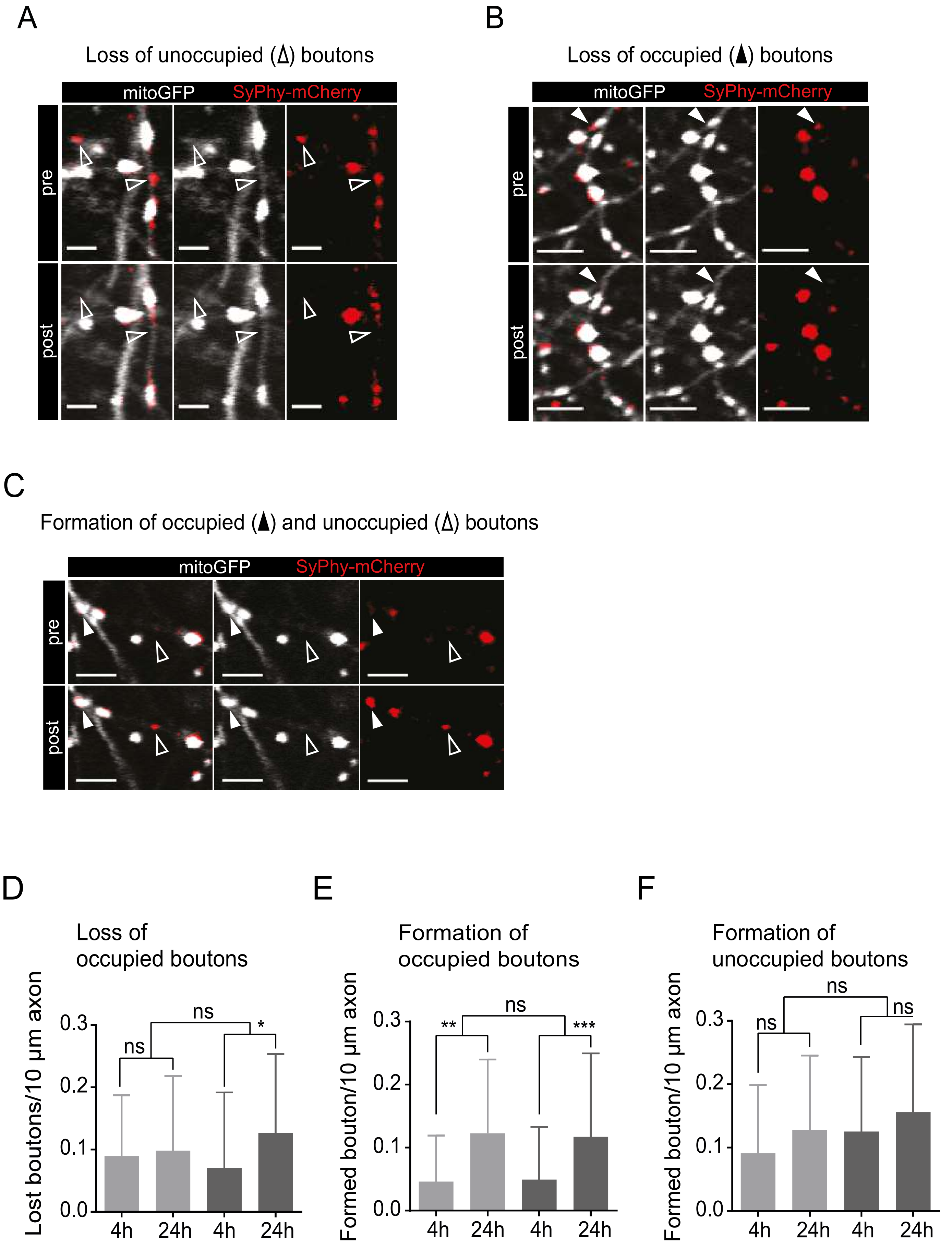


Supplementary Figure 5

### Supplemental Figure 6

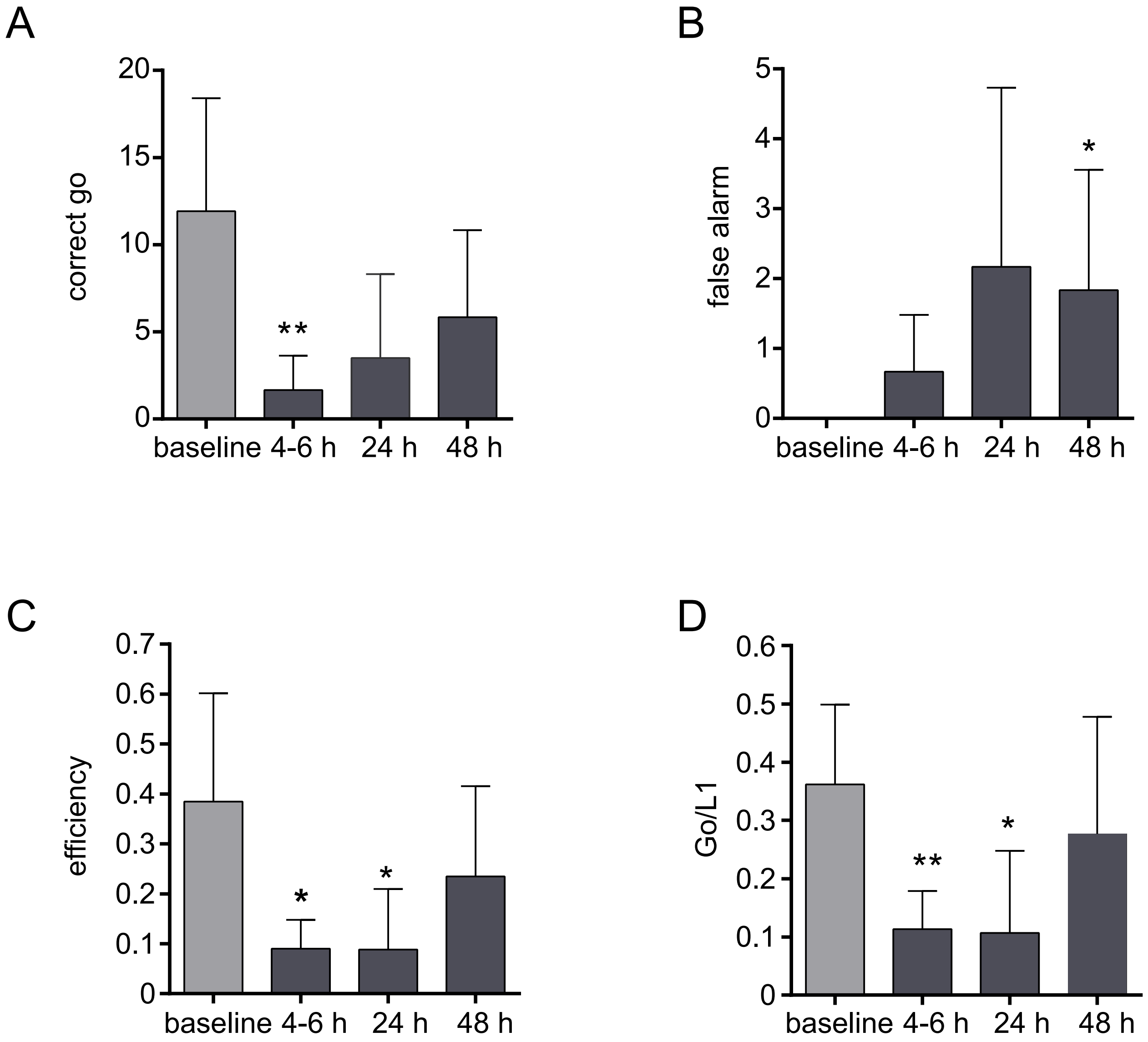


Supplementary Figure 6
