## Supplemental Table 2 for "Single-dose ethanol intoxication causes acute and lasting neuronal changes in the brain"

| Genotype | IA (1:36)* | EA + AA (1:36 + 1:400)* | Balance ** | n |
|---|---|---|---|---|
| TH-Gal4/+ | -0.50 ± 0.04 | -0.29 ± 0.02 | 0.08 ± 0.06 | 6 |
| UAS-milton-RNAi$^{GD}$/+ | -0.22 ± 0.07 | -0.37 ± 0.09 | 0.00 ± 0.05 | 6 |
| TH-Gal4/UAS-milton-RNAi$^{GD}$ | -0,37 ± 0.11 | -0.27 ± 0.08 | -0.10 ± 0.13 | 6 |
| TH-Gal4/+ | -0.45 ± 0.05 | -0.46 ± 0.07 | -0.03 ± 0.09 | 6 |
| UAS-milton-RNAi$^{TRiP}$/+ | -0.49 ± 0.11 | -0.60 ± 0.07 | -0.01 ± 0.03 | 6 |
| TH-Gal4/UAS-milton-RNAi$^{TRiP}$ | -0.58 ± 0.10 | -0.46 ± 0.13 | -0.11 ± 0.12 | 6 |
| UAS-dmiro-RNAi$^{TRiP}$/+ | -0.49 ± 0.07 | -0.35 ± 0.10 | -0.01 ± 0.04 | 6 |
| TH-Gal4/UAS-dmiro-RNAi$^{TRiP}$ | -0.46 ± 0.05 | -0.56 ± 0.10 | -0.07 ± 0.11 | 6 |

Supplementary Table 2
